# Comparative chromosome-scale genomics of the ayahuasca liana *Banisteriopsis caapi* reveals paleopolyploid ancestry and recent tandem-gene expansions as candidate signatures of ethnovariety differentiation

**DOI:** 10.64898/2026.10.11.758337

**Authors:** Vitor Trinca, Francisco Prosdocimi, Pedro Côrtes-Barros, Ana Paula Moraes, Mariana Cordeiro Almeida, Alexandre Rossi Paschoal, Saura Rodrigues da Silva, Vitor F. O. Miranda, Jose Beethoven Figueiredo Barbosa, Danilo Ribeiro de Oliveira, Simone Silva dos Santos Lopes, Alessandro M. Varani

## Abstract

**Background:** *Banisteriopsis caapi* is the Amazonian liana that provides the β-carboline-rich vine component of ayahuasca, a traditional Amazonian psychoactive preparation used in Indigenous and ceremonial contexts. The species encompasses culturally recognized ethnovarieties, some differing in stem architecture. Despite its cultural, ecological, and pharmacological importance, *B. caapi* remains largely unexplored at the genomic level. Here, we generated chromosome-scale genome assemblies for the caupuri and tucunacá ethnovarieties and used them to investigate genome evolution, structural variation, repeat composition, and candidate genomic features associated with ethnovariety-level differentiation.

**Results:** Caupuri and tucunacá assemblies span 1.16 and 1.13 Gb, respectively, each organized into ten high-quality chromosome-scale pseudomolecules. Both genomes retain signatures of two ancient post-gamma duplication components, with related signals also detected in *Malpighia emarginata*, suggesting that at least part of this paleopolyploid history may predate the divergence of these Malpighiaceae lineages. Against this conserved chromosomal framework, the sampled caupuri and tucunacá genomes differ in structural variation, repeat composition, and gene-family profiles. Transposable element-associated insertions and deletions dominate the structural-variant landscape, while both genomes are highly repetitive, with approximately 78% of their assemblies composed of repetitive DNA largely shaped by LTR retrotransposons, including *Gypsy*/Retand and *Copia*/Ale, *Copia*/Tork, and *Copia*/Ivana lineages, together with non-autonomous large retrotransposon derivatives (LARDs). Most genes belong to shared orthogroups, indicating a largely conserved gene repertoire. However, candidate ethnovariety-biased differences emerge from the combined remodeling of gene families, LARD and LTR-associated genomic neighborhoods, and comparatively recent local expansions of tandem-duplicated gene arrays encoding signaling receptors, transporters, hormone-associated genes, cell wall and secondary wall functions, and meristem or organ-development pathways.

**Conclusions:** These results indicate that the sampled caupuri and tucunacá genomes differ mainly through localized remodeling of shared genomic modules, including comparatively recent repeat turnover and local tandem-gene expansions. These assemblies provide the first large-scale nuclear genomic resource for *B. caapi* and a foundation for investigating paleopolyploid evolution in Malpighiaceae and candidate mechanisms of ethnovariety diversification.

## 1. Introduction

*Banisteriopsis caapi* (Spruce ex Griseb.) C.V.Morton is a Neotropical liana of the Malpighiaceae family, known as mariri or jagube, and occupies a central role in Amazonian ethnobotany as the principal liana component of ayahuasca, a preparation in which it is traditionally combined with *Psychotria viridis* Ruiz & Pav. leaves, known as chacrona [1–3]. Ayahuasca has a long-standing role in Indigenous and mestizo healing, ritual, and spiritual practices across the Amazon and has more recently expanded into biomedical, therapeutic, and other cultural contexts [1, 4, 5].

Recent phylogenetic and biogeographic analyses indicate a Miocene origin for *Banisteriopsis*, including the lineage leading to *B. caapi*, suggesting that its diversification happened more than 5 million years in the past, therefore predating largely human management [6]. The cultural and pharmacological relevance of ayahuasca has been extensively discussed, particularly because *B. caapi* provides β-carboline alkaloids that modulate the oral bioavailability of N,N-Dimethyltryptamine (DMT) and other tryptamine-based compounds present in the brew [7–9]. Large-scale chemical profiling of *B. caapi* and ayahuasca samples has revealed broad variation in harmine, harmaline, and tetrahydroharmine levels, indicating substantial phytochemical diversity within this species and its preparations [9].

Despite this broad ethnobotanical, pharmacological, and phytochemical interest, and despite the recent availability of a complete mitogenome for *B. caapi* [10], chromosome-scale nuclear genomic resources are still lacking, restricting conservation genetics managing, comparative genome analyses, repetitive DNA studies, gene-family evolution, and morphotype-level differentiation in this culturally significant and sacred Amazonian species. Placing *B. caapi* within a broader Malpighiales comparative framework is also relevant because well-developed genomic references are available for Salicaceae, including *Populus* and *Salix*, while *Malpighia emarginata* [11] provides a key comparative reference within Malpighiaceae.

Within *B. caapi*, Amazonian communities recognize distinct ethnovarieties that differ in their morphotypic characteristics, especially in the stem architecture. Caupuri develops prominent nodes along the liana, whereas tucunacá has a smoother stem surface (Figure 1) [1, 3, 12]. These ethnovarieties are associated with distinct ethnobotanical classifications and perceived properties in the ayahuasca brew [1, 3]. Previous DNA barcoding analyses in *Banisteriopsis* showed that nuclear markers, particularly ITS, recovered higher genetic diversity than plastid markers and identified lineages corresponding to the ethnovarieties used in ayahuasca preparation [12]. These observations raise the possibility that at least part of the ethnobotanical diversity recognized in *B. caapi* reflects biologically meaningful variation within the species. From an evolutionary genomics perspective, this system provides an opportunity to examine whether contrasting stem phenotypes are accompanied by differences in genome organization, structural variation, repetitive-element composition, and gene-family dynamics. Chromosome-scale comparative assemblies therefore provide a framework for evaluating whether morphotype-level traits are accompanied by copy-number changes, local duplications, and functional divergence among candidate genes and gene families involved in development, signaling, transport, and cell wall organization.

**Figure 1.**
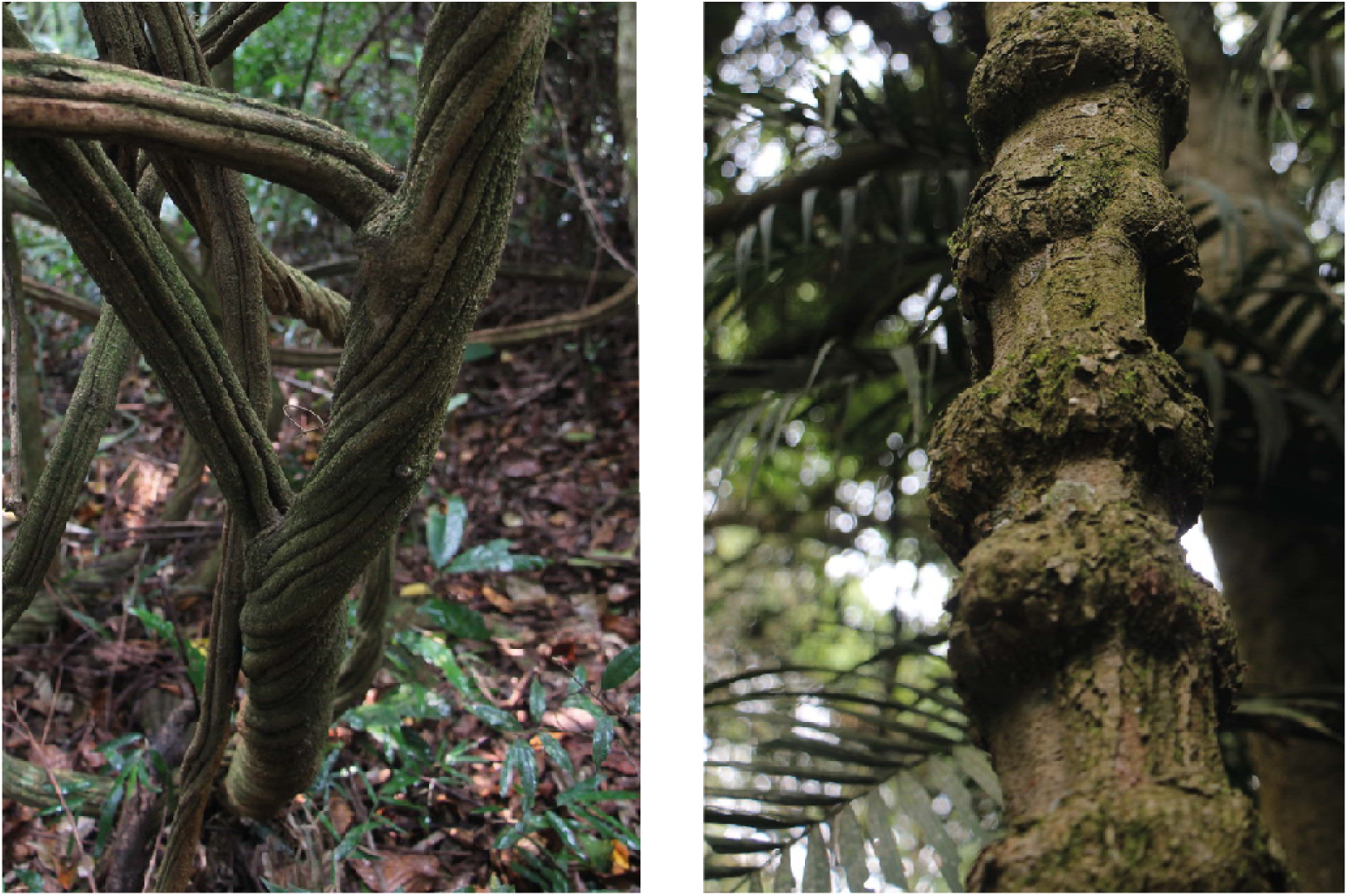
Detailed view of the two culturally recognized *Banisteriopsis caapi* ethnovarieties analyzed in this study. Tucunacá is shown on the left and caupuri on the right. Caupuri is characterized by prominent stem nodes, whereas tucunacá shows a smoother stem surface. In this study, these ethnovarieties were analyzed as morphologically distinct sampled accessions.

In plant genomes, morphotype-associated divergence can arise from multiple interacting sources of variation, including structural variants, transposable elements, whole-genome duplication, gene-family expansion and contraction, and recent local duplications [13, 14]. Repetitive elements are major contributors to genome size variation and can influence chromosomal organization, local recombination, nearby gene structure, and regulatory landscapes [15]. Likewise, ancient and recent duplication events provide raw material for functional diversification through differential gene retention, copy-number shifts, and lineage-biased expansion of specific gene families [14, 16]. Tandem duplication is particularly relevant in this context because it can generate local high-copy clusters involving genes associated with environmental response, signaling, specialized metabolism, and cell wall remodeling [14, 16]. Therefore, integrating chromosomal architecture, repeat landscapes, structural variation, and gene-family dynamics offers a powerful framework for identifying genomic features and candidate mechanisms associated with divergence between closely related morphotypes.

Here, we present chromosome-scale genome assemblies for the caupuri and tucunacá morphotypes of *B. caapi* and use them to investigate genome evolution and morphotype-level differentiation. We characterize assembly structure, gene content, repetitive-element composition, structural variation, centromeric and telomeric features, and signatures of post-γ paleopolyploid evolution. We further integrate orthogroup, gene-family, tandem-duplication, and functional-enrichment analyses to identify candidate genes, gene clusters, and gene families associated with differences between the sampled morphotypes. By integrating chromosome-scale comparative genomics this study provides the first chromosome-scale nuclear genomic resources for *B. caapi* and prioritizes candidate genomic features for future investigation of the morphological divergence between the caupuri and tucunacá ethnovarieties. It also provides a genomic foundation for future studies of conservation and population genetics aimed at assessing genetic diversity, guiding population management, and supporting strategies to mitigate the impacts of overexploitation on *B. caapi*.

## 2. Methods

### 2.1. Plant sampling, flow cytometry, nucleic acid extraction, and sequencing

Plant material corresponded to two culturally recognized ethnovarieties of *Banisteriopsis caapi*, caupuri and tucunacá, which were analyzed here as morphologically distinct sampled accessions. Ethnovariety identification followed the presence of prominent stem nodes in caupuri and their absence in tucunacá, together with the traditional knowledge of members of the Centro Espírita Beneficente União do Vegetal – Núcleo Menino Galante (CEBUDV, Serra da Cantareira, in the municipality of Mairiporã, Brazil; coordinates: 23.380562 S, 46.58857 W). Plant material was obtained from cultivated caupuri and tucunacá specimens maintained at the CEBUDV and deposited at the JABU Herbarium (São Paulo State University, Jaboticabal campus, Brazil) as vouchers JABU1322 and JABU1325. Leaves were dark-adapted for 24 h, flash-frozen in liquid nitrogen, and shipped to the Arizona Genomics Institute for high-molecular-weight DNA extraction and sequencing.

Genome size estimation followed standard flow cytometry protocol [17, 18] using ∼25 mg of leaf tissue macerated in the Ebihara buffer [19] with RNase (0.025 μg/mL). *Raphanus sativus* ‘Saxa’ was employed as an internal standard (2C = 1.11 pg [20]). Nuclei were stained with propidium iodide (1 mg/mL), analyzed on a FACSCanto II cytometer, and processed using FACSDiva and Flowing Software 2.5.1. Two individuals of caupuri and three individuals of tucunacá were analyzed in triplicate, retaining measurements with a coefficient of variation below 5% and a morphotype standard deviation below 3%. For statistical comparison of flow-cytometry genome-size estimates between morphotypes, technical triplicates were first averaged within each individual. Differences in individual mean 2C values between caupuri and tucunacá were then evaluated using Welch’s two-sample t-test.

High-molecular-weight DNA was extracted using a modified CTAB protocol [21], quantified using Qubit and NanoDrop, and inspected on Femto Pulse. Libraries were sheared to 10–30 kb using Covaris g-TUBEs and sequenced using PacBio HiFi technology. The caupuri library was sequenced on a single PacBio Revio SMRT Cell, whereas the tucunacá libraries were sequenced on two PacBio Sequel II SMRT Cells. GenomeScope 2.0 [22], Smudgeplot [22], and PloidyFrost [23] were used to estimate genome size, ploidy, and heterozygosity. Hi-C libraries were prepared only for caupuri using the Arima Genomics kit and sequenced on an Illumina NovaSeq platform (2 × 150 bp), with quality control performed using hic_qc scripts from Phase Genomics (commit: 6881c33; [24]). RNA was extracted from fresh leaves of both morphotypes using PureLink Plant RNA Reagent with cleanup using NucleoSpin, and only samples with RIN > 7 were sequenced. Long-read Iso-Seq libraries were sequenced on PacBio Sequel IIe, and short-read RNA-Seq libraries were sequenced on an Illumina HiSeq 2000 (2×100 bp).

### 2.2. Genome assembly and quality evaluation

PacBio HiFi reads from each ethnovariety were assembled independently with hifiasm v0.19.3 using default parameters [25]. Assembly statistics for the hifiasm primary assemblies were visualized using snail plots generated with BlobTK [26]. Primary assemblies were screened for potential contaminant sequences using Kraken 2 v2.1.3 [27], and non-plant-related contigs were removed before chromosome-scale scaffolding.

Hi-C scaffolding was performed using the caupuri Hi-C library. For tucunacá, caupuri-derived Hi-C reads were used only as cross-morphotype chromatin-contact information to order, orient, and join pre-assembled scaffolds into chromosome-scale pseudomolecules; they were not used to correct the hifiasm primary contigs. Hi-C read processing, contact-map generation, and chromosome-scale scaffolding were performed using the Juicer/3D-DNA workflow [28, 29], followed by manual inspection of scaffold ordering and orientation in Juicebox [30]. Final post-review scaffolds were generated with the 3D-DNA post-processing pipeline, and remaining scaffold gaps were refined using gap-closing procedures from the MaSuRCA package v4.1.1 [31]. Chromosome numbers were assigned primarily on scaffold length, while preserving the same numbering for corresponding chromosomes between the two morphotype assemblies.

Assembly completeness and accuracy were evaluated using complementary approaches. BUSCO v6 was run with the embryophyta_odb12 [32] dataset to estimate the recovery of conserved plant single-copy genes. Merqury v1.3 [33] was used for k-mer-based quality evaluation, including base-level consensus quality and k-mer completeness, whereas Inspector v1.0.2 [34] was used to assess read-to-assembly consistency, mapping rate, depth, and assembly error profiles. Repeat-space assembly quality was evaluated using the LTR Assembly Index (LAI) v2.9.8 [35]. Telomeric and centromeric repeated regions were identified using quarTeT (commit e1a2f72) [36] and the Centromics pipeline [37], respectively.

### 2.3. Transcriptome processing, gene prediction, and functional annotation

Iso-Seq and short-read RNA-seq data were generated from leaf tissue and used as qualitative transcript evidence for genome annotation. Iso-Seq reads were processed with SMRT Link 12.0 to generate high-confidence transcript sequences. Short-read RNA-seq data were assembled *de novo* with Trinity v2.14.0 [38] and aligned to each genome assembly with HISAT2 [39]. The high-confidence transcript sequences were aligned to the genome with minimap2 [40], and genome-guided transcript models were reconstructed with StringTie2 [41]. Iso-Seq transcripts, de novo transcript assemblies, and genome-guided transcript reconstructions were integrated with PASA v2.5.3 [42] and TransDecoder v5.7.0 [43] to generate transcript structures used as evidence for structural annotation. Transcriptome completeness was evaluated with BUSCO v6 using the embryophyta_odb12 dataset.

Protein-coding gene prediction was performed on the repeat-masked (soft-masked) genome assemblies by integrating *ab initio*, transcript-supported, and protein homology-based evidence. Gene prediction evidence included BRAKER3 [44], PASA transcript models, EVidenceModeler [42] consensus models, and protein-based alignments generated from plant reference proteins from embryophyta_odb12 dataset. The annotation workflow followed the plant genome annotation strategy previously applied by our group, which benchmarks and integrates multiple evidence sources to reduce unsupported gene models [45–48]. Predicted gene models overlapping structurally annotated transposable elements or containing TE-associated domains were inspected during post-processing and removed when they lacked transcript, protein, or conserved-domain support. The final gene sets were evaluated with BUSCO in protein mode.

Functional annotation was assigned by integrating similarity- and domain-based evidence in Blast2GO. Predicted proteins were compared against plant protein databases using BLAST-based searches [49], and complementary functional evidence from eggNOG-mapper [50] and InterProScan [51] was incorporated into the Blast2GO annotation framework [52]. Gene descriptions, COG categories [53], Enzyme Commission numbers [54], Gene Ontology (GO) terms [55], Pfam domains [56], InterPro domains [57], and additional annotation fields were exported from the integrated annotation output and compiled into the final gene annotation tables. Gene-duplication topology classes, including singleton, dispersed, proximal, tandem, transposed, and WGD-derived genes, were assigned using MCScanX [58] and DupGen_finder [14] based on chromosome-anchored gene coordinates, homologous gene-pair relationships, and syntenic collinearity, and were added to the final annotation tables. Descriptive Transcripts Per Million (TPM) values were estimated from genome-guided RNA-seq alignments and used only as annotation support. For comparative analyses, the *M. emarginata* genome was reannotated using the same annotation workflow.

### 2.4. Prediction and analysis of non-coding RNAs

Non-coding RNA (ncRNA) loci were predicted using complementary sequence-similarity covariance-model, and specialized tRNA-search strategies. BLAST tools [49] were used to search against all ncRNAs from the RNA central database release 26 [59] and the microRNAs from the curated Plant microRNA Gene Database, PmiREN 3.0 [60], in accordance with current best practices and computational guidelines for miRNA annotation [61]. Transfer RNA loci were annotated independently with tRNAscan-SE (v.2.0.9 July 2021) [62]. Predictions flagged by tRNAscan-SE as possible pseudogenes were retained in the final dataset and identified accordingly. For RNAcentral searches, an E-value threshold of 1 × 10 and minimum identity and coverage of 90% were applied, whereas minimum identity and coverage of 80% were used for miRNA searches. Structural ncRNA searches were performed with INFERNAL v.1.1.3 [63] against the Rfam 15.1 (January 2026, 4,227 families) database [64]. The INFERNAL annotation used cmscan software with the parameter --cut_ga --rfam –nohmmonly --clanin Rfam.clanin --oskip. Filtering and merging were performed manually, retaining only eukaryotic predictions located on chromosome-level sequences.

### 2.5. Repeat annotation, repeat landscapes, TE-density, and LARD–gene associations

Repetitive elements were annotated using EDTA-GUI plant repeat annotation workflow [65] applied independently to the caupuri, tucunacá, and *M. emarginata* genome assemblies. Repeat divergence landscapes were generated from the EDTA-GUI annotation outputs by summarizing Kimura divergence values across repeat classes. Intact LTR insertion-time distributions were estimated separately for LTR/*Gypsy* and LTR/*Copia* elements using the pairwise divergence between the two LTRs of each intact element. Insertion ages were standardized using a neutral substitution rate of 1.1 × 10 substitutions per site per year, following the rate adopted for *B. caapi* in the EDTA-GUI reference database [66].

TE-density profiles around annotated genes were calculated with TE_Density[67]. The final comparative plots included caupuri, tucunacá, and *M. emarginata*, using the same TE categories and a window series ranging from 1.5 kb to 15 kb, with 1 kb increments between consecutive windows.

Associations between LARD elements and annotated genes were evaluated through coordinate-based intersection analyses implemented with custom scripts. Each gene–LARD pair was assigned to one of four mutually exclusive positional categories: gene_inside_LARD, when the complete gene interval was contained within a LARD annotation; LARD_inside_gene, when the complete LARD interval was contained within the gene span; partial_overlap, when the two intervals overlapped without reciprocal full containment; and flank-only, when a LARD element was located within 5 kb upstream or downstream of a gene without overlapping the gene body. For genes associated with multiple LARD elements or multiple positional categories, a single representative relationship was retained using a hierarchical classification scheme in the following order: gene_inside_LARD, LARD_inside_gene, partial_overlap, and flank-only. The resulting gene-level LARD-association matrix was integrated with functional annotations, orthogroup assignments, and duplication-topology classifications.

Functional enrichment analyses were performed independently for caupuri and tucunacá. Genes classified as gene_inside_LARD, LARD_inside_gene, or partial_overlap were used as the tested LARD-associated gene set, whereas genes classified only as flank-only were excluded from these tests. For each genome, the background consisted of all chromosome-anchored genes with functional annotation or duplication-topology classification. Gene Ontology enrichment was performed with GOATOOLS [68], using term propagation to parental GO categories, one-sided Fisher’s exact tests, and Benjamini–Hochberg false-discovery-rate correction. Enrichment of COG categories, Enzyme Commission annotations, Pfam domains, and duplication-topology classes was evaluated separately using one-sided Fisher’s exact tests followed by Benjamini– Hochberg correction. Enriched categories were retained when they included at least three LARD-associated genes, showed fold enrichment greater than 1, and had FDR ≤ 0.05. Functional and duplication-topology profiles were also compared between caupuri and tucunacá using two-sided Fisher’s exact tests followed by Benjamini–Hochberg correction. *Malpighia emarginata* was not included in the enrichment analyses because only 17 non-flanking LARD-associated genes were recovered.

### 2.6. Synteny, whole-genome duplication analyses, and ancestral karyotype reconstruction

Comparative synteny analyses were performed using the chromosome-scale assemblies and predicted protein-coding genes of caupuri, tucunacá, *M. emarginata*, *Populus trichocarpa*, *Salix purpurea*, and *Vitis vinifera*. For each genome, the longest representative protein sequence per gene was used when multiple isoforms were present. Pairwise protein similarity searches were performed prior to collinearity detection, and syntenic blocks were inferred using MCScanX [58], DupGen_finder [14], WGDI [69], and the Python implementation of MCScan [70]. The resulting collinearity files were used to estimate syntenic depth among genomes and to generate chromosome-scale synteny visualizations.

Whole-genome duplication signatures were investigated from synonymous substitution rate distributions of duplicated gene pairs. For each set of collinear paralogous pairs, protein sequences were aligned with MAFFT [71], converted to codon alignments with PAL2NAL [72], and used to estimate synonymous divergence values with KaKs_Calculator v3.0 [73]. Ks distributions were summarized using all syntenic pairs, block-average estimates, and block-median estimates. Density plots were generated for the complete Ks range and for selected intervals used to inspect individual duplication components.

Additional WGD analyses were performed with WGD_v2 v2.0.38 [74]. Anchor-pair Ks distributions were used as input for Gaussian mixture modeling to identify major duplication components in each genome. Approximate duplication-time distributions were then estimated from Ks-derived intervals using the substitution-rate framework implemented in the WGD_v2 workflow. These time estimates were interpreted cautiously because Ks-based dating is sensitive to substitution-rate assumptions and rate heterogeneity among lineages. These analyses were performed independently for the caupuri and tucunacá genomes and repeated for comparative genomes used to contextualize the duplication signals. In the *M. emarginata*-focused WGD_v2 analysis, the two species anchor-pair components were represented as pseudo-taxa, whereas caupuri and tucunacá were retained as separate terminal taxa in the MCMCTree [75] chronogram. The posterior age recovered for the caupuri–tucunacá node was used as an indirect temporal estimate.

To examine chromosome-level consequences of ancient genome duplications, ancestral eudicot karyotype blocks were reconstructed using the ancestral karyotype reconstruction framework implemented in WGDI [69] and projected onto the chromosomes of caupuri, tucunacá, *M. emarginata*, *P. trichocarpa*, and *S. purpurea*. The WGDI-derived ancestral eudicots karyotype (AEK) background was used to assign retained genomic segments to ancestral protochromosomes and to visualize their distribution across modern chromosomes. Retained gene counts were summarized for each ancestral block and chromosome combination. The genome assemblies used for comparative and BUSCO-based phylogenetic analyses are listed in Table S19.

### 2.7. Structural variation analysis

Structural variants between the caupuri and tucunacá assemblies were identified from reciprocal whole-genome alignments using MUM&Co v3.8 [76], based on MUMmer4 v.4.0.0 [77] alignments. Chromosome-scale scaffolds were used as the primary input, and structural variants shorter than 50 bp were excluded from downstream summaries. The two morphotypes were alternately used as reference and query to recover candidate genome-specific insertions and deletions while reducing reference-direction bias.

Detected variants were classified into deletions, insertions, duplications, inversions, contractions, and translocations according to the MUM&Co output classes. To distinguish TE-associated events from variants without direct overlap with annotated repetitive elements, insertion and deletion intervals were intersected with the corresponding TE annotation files. Insertions or deletions overlapping annotated TEs were classified as mobile element-associated events, whereas those without TE overlap were retained as novel insertions or deletions.

Structural variants were summarized by chromosome, event class, frequency, and cumulative genomic span. For genome-coverage estimates, translocation spans were reported separately because large-scale alignment breaks and rearranged regions can inflate cumulative span estimates. Chromosome-wise SV tables were generated for both morphotypes and used to summarize total event counts and affected genomic space.

### 2.8. Orthogroup inference and gene family evolution

Because the complete caupuri and tucunacá annotations differed in the number of genes located on unanchored scaffolds, the primary orthogroup and gene-family analyses for *B. caapi* were performed using only genes located on chromosome-scale scaffolds. Orthogroups (OGs) were inferred with OrthoFinder v2.5.5 [78] using the comparative protein sets listed in Table S19, and the final *B. caapi* gene-family matrix retained only genes located on chromosome-scale scaffolds.

Genes classified as unassigned by OrthoFinder were not included in the CAFE5 [79] input matrix. Before CAFE5 analysis, families present in only one species and families with ≥100 copies in at least one species were removed to reduce the influence of lineage-specific artifacts and extreme copy-number variation, following the CAFE5 recommendation [79]. The species tree was converted to an ultrametric-time tree using TimeTree5-based [80] divergence constraints. Because caupuri and tucunacá are accessions of *B. caapi* rather than independent species, their split was treated as a short operational within-species branch for numerical stability and was not interpreted as a species-level divergence. A global error model was estimated from the filtered gene-family count matrix and applied to the final CAFE5 run. Significant gene-family shifts were inferred from the CAFE5 family-wide and branch-specific statistical outputs using P < 0.05. Functional annotations, GO terms, Pfam domains, duplication topology, and representative gene descriptions were integrated with the CAFE5 output to characterize expanded and contracted gene families in the two *B. caapi* morphotypes.

### 2.9. Prioritization of candidate orthogroups and clusters associated with differences between sampled morphotypes

A focused prioritization of candidate OGs potentially associated with differences between the sampled caupuri and tucunacá genomes was performed from significant CAFE5 orthogroups. Candidate prioritization was based primarily on GO evidence associated with plant architecture, meristem and organ development, shoot or stem branching, hormone response, transport and trafficking, signaling receptors, cell wall organization, secondary wall formation, vascular differentiation, transcriptional regulation, and related anatomical terms. Pfam and InterPro domains were used as supporting evidence, and annotation keywords were used only as fallback evidence when GO or domain information was incomplete. Each OG was assigned to one or more candidate functional categories and given a priority score based on the combined evidence. This prioritization was used as a hypothesis-generating procedure and was not interpreted as evidence of causal involvement in stem-architecture divergence.

As a complementary pairwise gene-cluster analysis, the chromosome-anchored caupuri and tucunacá protein sets used in the primary orthogroup analyses were compared with OrthoVenn3 [81]. Cluster membership was obtained from the OrthoVenn3 orthogroups output and classified as shared, caupuri-only, or tucunacá-only according to species composition. The ortholog, co-ortholog, and in-paralog outputs were used to audit the relationship structure, while cluster definitions were retained from the orthogroups output. Genes from exclusive clusters were integrated with gene-level functional annotations, OrthoFinder orthogroup assignments, reconstructed terminal CAFE5 events, and prioritized CAFE5 candidate orthogroups. Shared OrthoVenn3 clusters assigned to the same OrthoFinder orthogroups were also recorded to determine whether an exclusive protein cluster occurred within a broader gene family represented in both genomes. CAFE5 support was considered directionally concordant when a caupuri-only cluster was associated with an orthogroup expanded in caupuri and/or contracted in tucunacá, or when a tucunacá-only cluster was associated with an orthogroup expanded in tucunacá and/or contracted in caupuri. Clusters dominated by TE-related annotations, defined as at least 50% of the annotated genes, and clusters lacking informative gene descriptions were excluded only from the integrated candidate subset.

### 2.10. Tandem duplication analysis and functional enrichment of tandem-duplicated genes

Tandem-duplicated genes were identified from the duplication-topology classification and grouped into tandem arrays according to their chromosome location, physical proximity, and consecutive order. Adjacent tandem pairs were defined as consecutive gene pairs within the same tandem array. For each array, chromosome, start and end coordinates, array size, span, strand composition, orthogroup composition, and CAFE5 status were summarized. Adjacent tandem pairs were then intersected with the prioritized candidate orthogroups. A pair was considered candidate-related when at least one gene belonged to a prioritized candidate OG potentially associated with differences between the sampled genomes. Candidate categories associated with tandem pairs were inherited from the corresponding candidate OG. The final tandem table reports candidate-related adjacent tandem pairs and their associated annotations, orthogroups, CAFE5 status, and available Ka/Ks fields.

Functional enrichment of tandem-duplicated genes was performed independently for the caupuri and tucunacá genomes using the final gene annotation tables and duplication-topology classifications. For each morphotype, genes classified as tandem duplicated were used as the study set, whereas all genes from the same genome with at least one valid functional annotation were used as the corresponding population set. GO enrichment was performed with GOATOOLS, Fisher’s exact test, propagation of counts to parental GO terms, and Benjamini– Hochberg false-discovery-rate correction (FDR). Enriched GO terms with FDR ≤ 0.05 were retained for downstream interpretation. Complementary enrichment analyses were performed for COG functional categories and Enzyme Commission (EC) annotations using custom Python scripts. For COG and EC analyses, genes with at least one COG category or EC number were used as annotation-specific population sets, and tandem genes with the corresponding annotation were used as study sets. For each COG category or EC number, overrepresentation in the tandem set was tested using one-sided Fisher’s exact tests followed by Benjamini– Hochberg FDR correction. Terms with FDR ≤ 0.05 were retained as significant.

To evaluate whether tandem-duplicated genes showed differences in functional annotation profiles between the sampled genomes, additional pairwise comparisons were performed between the complete GO-, COG-, and EC-annotated gene sets of caupuri and tucunacá, and between their tandem-duplicated gene sets. For these between-morphotype comparisons, terms represented by at least three genes in either group were tested using Fisher’s exact tests, followed by Benjamini–Hochberg correction. The resulting enriched terms were used to distinguish annotation signatures associated with tandem duplication within each genome from broader differences in functional annotation composition between the two sampled genomes.

## 3. Results

The genomes of both varieties were assembled into approximately 1.1 Gb haploid genomes, each comprising ten chromosomes, with minimal gaps and more than 99% of genes successfully predicted (Table 1), as detailed below.

**Table 1.**
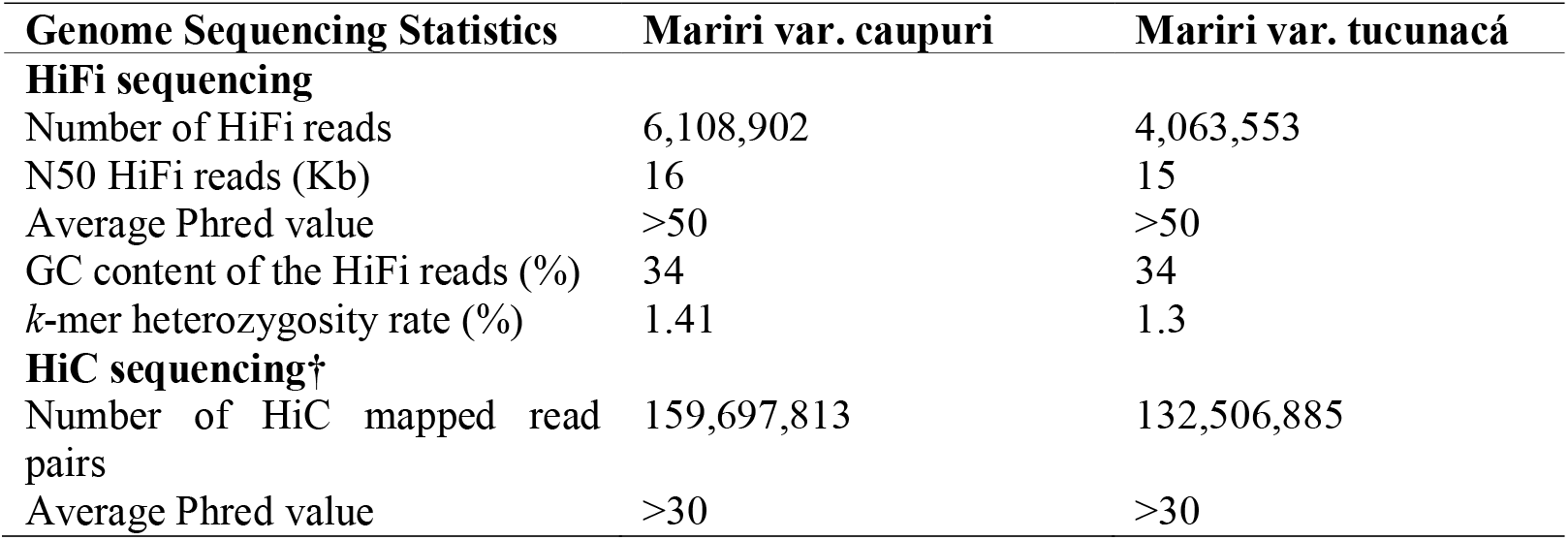
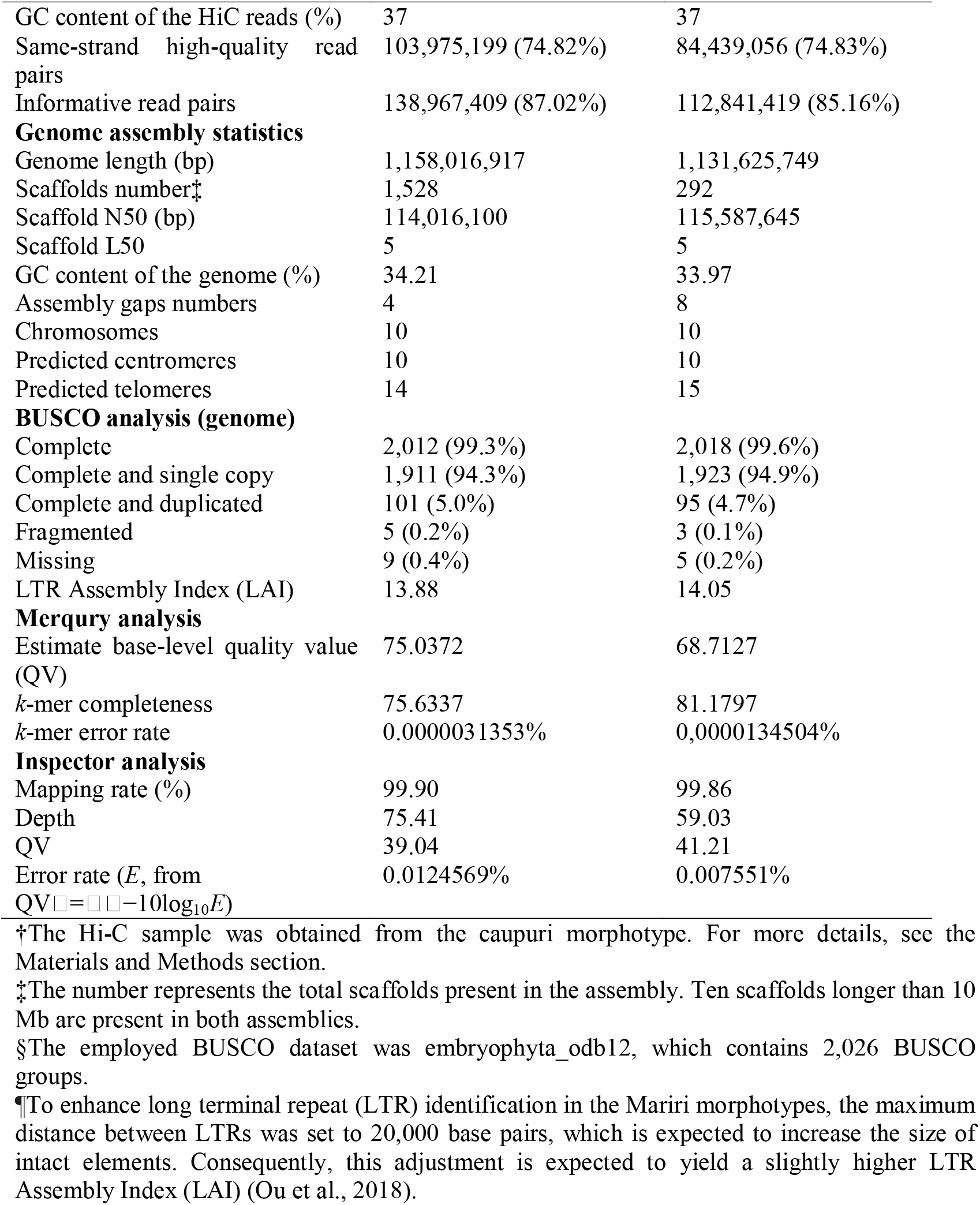
Statistics of *Banisteriopsis caapi* (mariri) genome sequencing and assembly for the two different morphotypes, caupuri and tucunacá.

### 3.1. DNA Sequencing and Assembly

Genome size estimates obtained by flow cytometry indicated highly similar nuclear DNA contents in the sampled *Banisteriopsis caapi* morphotypes (Table S1). Caupuri exhibited a 2C value of 2.66 pg (± 0.07, CV = 4.0; 1C = 1.30 Gb), while tucunacá showed 2C = 2.70 pg (± 0.046, CV = 3.91; 1C = 1.32 Gb). A Welch two-sample t-test performed on individual mean 2C estimates did not detect a significant difference between the sampled morphotypes (t = −0.684, df = 1.10, p = 0.609). Long-read PacBio sequencing generated 6.1 million reads for caupuri (N50 = 16 kb; one Revio SMRT Cell) and 4.0 million reads for tucunacá (N50 = 15 kb; two Sequel II SMRT Cells) (Table S1). K-mer–based analyses supported a diploid genomic profile for both morphotypes (Figure S1). GenomeScope estimated genome sizes of 1.06 Gb for caupuri and 1.08 Gb for tucunacá, corresponding to ∼81% of the cytometric 1C estimates (Table S1). The heterozygous k-mer components were estimated at 1.4% in caupuri and 1.3% in tucunacá. These k-mer-derived genome-size estimates were lower than those obtained by flow cytometry but close to the final assembled genome sizes. Sequencing depth was high for both morphotypes, ∼90× for caupuri and ∼60× for tucunacá. Hi-C sequencing was performed for the caupuri morphotype. The caupuri Hi-C data were used to scaffold the caupuri assembly and, for tucunacá, were used only as cross-morphotype chromatin-contact information to order, orient, and join pre-assembled scaffolds into chromosome-scale pseudomolecules (Table S2).

The primary HiFi assemblies were highly contiguous and contained almost all ten assembled chromosomes (Figure S2). The draft assembly combined with Hi-C-assisted scaffolding and cross-morphotype chromatin-contact support produced highly contiguous chromosome-scale genome assemblies (Table 1; Figure 2A; Table S2). The caupuri assembly spans 1.16 Gb across 1,528 scaffolds, with an N50 of 114.0 Mb, L50 of 5, and only four annotated gaps. Its ten chromosome-scale scaffolds cover 1.10 Gb (95% of the total assembly). Tucunacá yielded a similarly contiguous 1.13 Gb assembly comprising 292 scaffolds, with an N50 of 115.6 Mb, L50 of 5, and eight annotated gaps. Chromosome-scale scaffolds represent 1.11 Gb (98% of the total) (Table 1). Centromeric repeats were identified on all chromosome-level scaffolds in both morphotypes (Table S3). Telomeric repeats occurred at both ends of four caupuri chromosomes and at one end of the remaining six; in tucunacá, five chromosomes exhibited complete telomeric boundaries, and five had telomeric motifs at a single end (Table S4).

**Figure 2.**
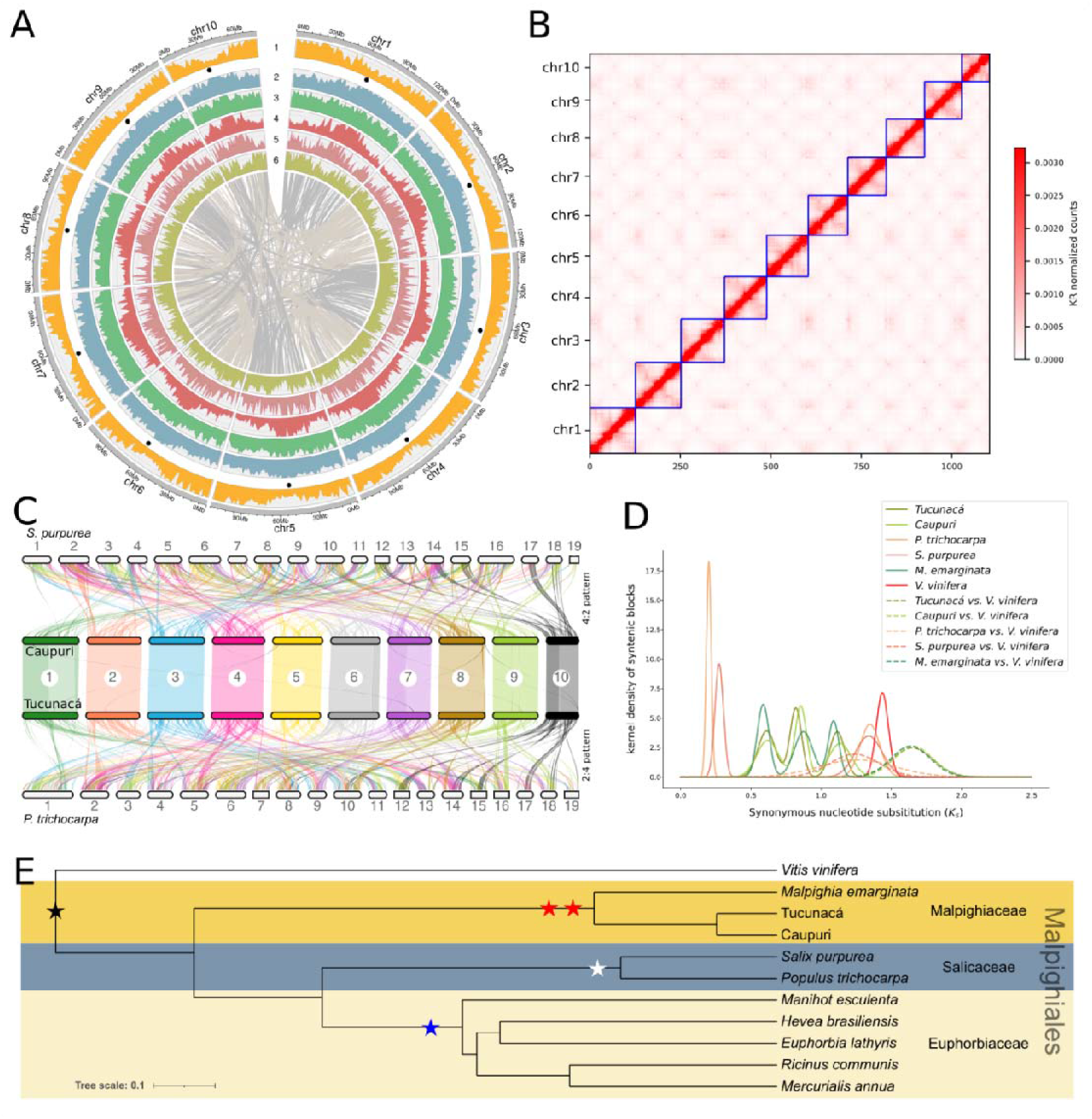
Summary features of *Banisteriopsis caapi* genome assembly. (**A**) Circular genomic plot of *Banisteriopsis caapi*, caupuri morphotype. From the outer to the inner circles: (1) genes, and the black dots represent the predicted centromeric positions. (2) LTR *Copia*, (3) LTR *Gypsy*, (4) LINE, (5) TIR, and (6) Helitron. (**B**) Caupuri morphotype Hi-C contact map; Tucunacá morphotype genomic plot and Hi-C contact maps are shown in Figure S3. (**C**) Synteny blocks comparison between the caupuri and tucunacá morphotypes, *S. purpurea*, and *P. trichocarpa*. This denotes a 2:4 genome duplication pattern between *B. caapi* and the Salicaceae species. (**D**) WGD_v2 analysis of anchor-pair Ks distributions in caupuri and tucunacá. Two intermediate duplication components were recovered in both genomes. Approximate duplication-time estimates for anchor pairs within the Ks interval of 0.51–2.11 ranged from 1.72 to 28.39 Mya, with median estimates of 8.78 Mya in caupuri and 7.75 Mya in tucunacá. These distributions provide temporal context for the shared paleopolyploid signals and do not represent the age of a single duplication event. Solid lines represent Gaussian-fitted distributions from within-species syntenic homologous pairs, whereas dashed lines represent interspecific distributions between each respective species and *V. vinifera* (**E**) BUSCO phylogenetic tree for the sampled Malpighiales and *V. vinifera* as the outgroup species. Stars represent the identified Mariri WGDs (red star), the confirmed Salicaceae WGD (white star), and the Euphorbiaceae WGD event (blue star) [82]. Black star represents the γ triplication [83].

Assembly completeness metrics further demonstrate the high quality of both genomes. BUSCO analyses recovered 99.3% of expected conserved genes in caupuri and 99.6% in tucunacá. LTR Assembly Index (LAI) scores reached 13.88 and 14.04, respectively, supporting reference-quality assembly of LTR-rich repetitive regions. Independent evaluations using Merqury and Inspector corroborated the high consensus accuracy, low error rates, and high k-mer completeness of both assemblies (Table 1).

### 3.2. Structural annotation and chromosome-anchored gene content

Structural annotation predicted 59,836 genes in the complete caupuri assembly and 47,458 genes in the complete tucunacá assembly, although this difference was largely associated with genes located on unanchored scaffolds (Table S2; Table S5; Figure S3). However, since caupuri and tucunacá have only 1 and 2 BUSCO groups classified as complete in the haplotig-predicted genes, only the genes predicted on chromosome-length scaffolds were retained for further analysis. Caupuri contained 42,608 genes and 64,497 CDSs, including 64,363 complete CDSs, whereas tucunacá contained 42,486 genes and 66,537 CDSs, including 66,392 complete CDSs. The annotated gene space covered 11.70% of the caupuri assembly and 11.16% of the tucunacá assembly (Table 2). Annotation completeness was high in both morphotypes, with BUSCO recovering 98.8% complete genes in caupuri and 99% in tucunacá. Complete duplicated BUSCOs represented around 4% in both annotations (Table 2; Table S2). OMArk assigned 86.01% of predicted genes in caupuri and 85.09% in tucunacá to taxonomically consistent Rosid orthogroups (Table S2). Complete gene-level structural and functional annotations for the chromosome-anchored gene sets are provided in Table S6.

**Table 2.** Banisteriopsis caapi (Mariri) and Malpighia emarginata annotation features.

| Features | *B. caapi*<br>var. caupuri | *B. caapi*<br>var. tucunacá | *M. emarginata* |
| --- | --- | --- | --- |
| **Repetitive sequence annotation** |  |  |  |
| Class I elements | 55.47% | 57.24% | 67.96% |
| *LTR/Gypsy* | 28.39 % | 28.33% | 38.48% |
| *LTR/Copia* | 13.30% | 15.07% | 17.06% |
| LTR non-autonomous | 10.43% | 9.52% | 9.65% |
| Non-LTR | 2.09% | 3.27% | 2.41% |
| Class II elements | 7.69% | 8.28% | 21.83% |
| TIRs | 6.08% | 6.47% | 20.09% |
| Helitron | 1.61% | 1.80% | 1.74% |
| Unclassified | 15.82% | 14.09% | 3.34% |
| **Genome annotation†** |  |  |  |
| Number of genes | 42,608 | 42,486 | 36,031 |
| Number of CDSs (including isoforms) | 64,497 | 66,537 | 57,845 |
| Complete CDS | 64,363 | 66,392 | 57,651 |
| Start, no stop CDS | 46 | 57 | 61 |
| Stop, no start CDS | 56 | 55 | 79 |
| No stop, no start CDS | 32 | 33 | 33 |
| Mean gene length | 3,101 | 2,995 | 3,372 |
| Mean CDS length | 1,282 | 1,295 | 1,251 |
| Mean exons per gene | 5 | 5 | 6 |
| Mean introns per gene | 4 | 4 | 5 |
| % of genome covered by genes | 11.7 | 11.16 | 11.55% |
| % of genome covered by CDS | 4.29 | 4.22 | 6.88% |
| **Non-coding RNA annotation** |  |  |  |
| Total ncRNA loci | 6,213 | 6,345 | 5,463 |
| tRNA ‡ | 3,750 | 3,894 | 1,188 |
| snoRNA | 1,201 | 1,208 | 2,017 |
| rRNA | 512 | 468 | 1,745 |
| snRNA | 308 | 323 | 163 |
| miRNA | 172 | 170 | 162 |
| SRP_RNA | 29 | 27 | 28 |
| Ribozyme | 1 | 1 | 3 |
| RNase_RNA | 1 | 1 | 2 |
| lncRNA | 0 | 0 | 4 |
| **BUSCO analysis (annotation)§** |  |  |  |
| Complete | 2,001 (98.8%) | 2,005 (99.0%) | 2,008 (99.1%) |
| Complete and single copy | 1,913 (94.4%) | 1,918 (94.7%) | 1,098 (54.2%) |
| Complete and duplicated | 88 (4.3%) | 87 (4.3%) | 910 (44.9%) |
| Fragmented | 15 (0.7%) | 10 (0.5%) | 8 (0.4%) |
| Missing | 10 (0.5%) | 11 (0.5%) | 10 (0.5%) |

† Genome annotation for the chromosome-length scaffolds; for the complete genome annotation, see Table S5 and Table S6.
‡ Include predictions flagged as possible pseudogenes.
§ The employed BUSCO dataset was *embryophyta\_odb12*, which contains 2,026 BUSCO groups.

### 3.3. Genome-wide Identification and Annotation of Non-Coding RNAs

To delineate the ncRNA landscape across the three Malpighiaceae genome assemblies (caupuri, tucunacá, and *M. emarginata*), a comprehensive genome-wide annotation was performed. A total of 18,021 eukaryotic ncRNA loci were identified across the three assemblies (Table 2 and Table S7). The annotated ncRNA complement comprised nine major structural and functional classes: transfer RNAs (tRNAs), small nucleolar RNAs (snoRNAs), ribosomal RNAs (rRNAs), small nuclear RNAs (snRNAs), microRNAs (miRNAs), signal recognition particle RNAs (SRP_RNAs), ribozymes, RNase RNAs (RNase P/MRP), and long non-coding RNAs (lncRNAs). Comparative analysis revealed similar total ncRNA counts and class composition between the caupuri (6,213 loci) and tucunacá (6,345 loci) genomes. Both assemblies were dominated by tRNAs, which constituted 60.4% (3,750 loci) and 61.4% (3,894 loci) of their total ncRNA complements, respectively. Of these tRNA predictions, 738 in caupuri and 696 in tucunacá were flagged by tRNAscan-SE as possible pseudogenes. The high tRNA counts therefore partly reflect the retention of predicted pseudogene or repetitive tRNA-derived loci in the chromosome-scale assemblies (Table 2 and Table S7)

In contrast, *M. emarginata* exhibited a distinct ncRNA annotation profile, with 5,463 predicted loci and lower representation of tRNA-like loci (21.7%; 1,188 loci), but larger numbers of rRNA (31.9%; 1,745 loci) and snoRNA predictions (36.9%; 2,017 loci). Because rRNA genes occur in highly repetitive tandem arrays and plant snoRNAs may be organized in duplicated or polycistronic clusters, these differences may reflect variation in repetitive-locus representation, copy number, and annotation sensitivity. They were therefore not interpreted as evidence of lineage-specific functional expansion (Table 2 and Table S7).

Across all three genomes, several lower-abundance non-coding RNA classes showed broadly similar counts. miRNAs ranged from 162 to 172 loci, whereas snRNAs involved in spliceosomal machinery ranged from 163 to 323 loci. Signal recognition particle RNAs (27–29 loci), ribozymes (1–3 loci), and RNase P/MRP RNAs (1–2 loci) showed similar low counts across the three assemblies. Finally, despite recent advances in plant genome assembly, accurately annotating and functionally characterizing lncRNAs in non-model species still poses a significant bioinformatic challenge [84].

### 3.4. Paleopolyploidy and syntenic duplication signals

Comparative synteny analysis between the *B. caapi* genomes and those of *Populus trichocarpa* and *Salix purpurea* revealed fragmented yet detectable collinear relationships, consistent with the different chromosome numbers and lineage-specific chromosomal rearrangements that separate *B. caapi* from Salicaceae. Despite this fragmentation, syntenic depth patterns supported an approximate 2:4 relationship between *B. caapi* and the Salicaceae genomes (Figure 2C). Syntenic blocks were distributed across the ten *B. caapi* chromosomes and the 19 chromosomes of *P. trichocarpa* and *S. purpurea*, providing a comparative framework for evaluating ancient duplication patterns in *B. caapi*.

Ks-based analyses of syntenic paralogous and orthologous gene pairs using the WGDI pipeline [69] recovered multiple duplication-related peaks across the sampled genomes (Figure 2D). A lower-Ks peak around 0.33 was detected in the Salicaceae comparisons, whereas intermediate peaks around Ks ≈ 0.9 and Ks ≈ 1.2 were detected in the *B. caapi* genomes. A broader high-Ks component around Ks ≈ 1.1–1.4 was also recovered across the sampled species. WGD_v2 analyses [74] produced Ks distributions comparable to those obtained with WGDI and supported two intermediate duplication components in both *B. caapi* genomes, centered around Ks ≈ 0.8 and Ks ≈ 1.4 (Figure 2D; Figure S4; Figure S5). Approximate duplication-time estimates for anchor pairs within the Ks interval of 0.51–2.11 ranged from 1.72 to 28.39 Mya, with median estimates of 8.78 Mya in caupuri and 7.75 Mya in tucunacá (Figure S5). The broadly overlapping distributions were consistent with the shared duplication components recovered in the two genomes. Additional WGD_v2 analyses in *M. emarginata* recovered comparable intermediate Ks components, although with older apparent time estimates for the most recent component, ranging from approximately 28 to 50 Mya and showing a median of 40.61 Mya (Figure 2D; Figure S6). The same *M. emarginata*-focused analysis recovered an indirect posterior mean age of 1.13 Ma for the caupuri–tucunacá node, with a 95% highest-posterior-density interval of 0.31–2.26 Ma.

Ancestral eudicot karyotype reconstruction assigned retained AEK-derived genomic segments to chromosomes of *B. caapi*, *M. emarginata*, *P. trichocarpa*, and *S. purpurea* (Figure 3). The reconstruction included seven ancestral protochromosomes and 23,367 retained ancestral genes. Retained gene counts were summarized for the *B. caapi* genomes, *M. emarginata*, *P. trichocarpa*, and *S. purpurea*, showing the distribution of ancestral blocks across the modern chromosomes of each genome. Together, these syntenic, Ks-based, and ancestral-karyotype analyses support the presence of two ancient post-γ duplication components in *B. caapi*, embedded within a broader paleopolyploid background that may extend to other Malpighiaceae lineages (Figure 3).

**Figure 3.**
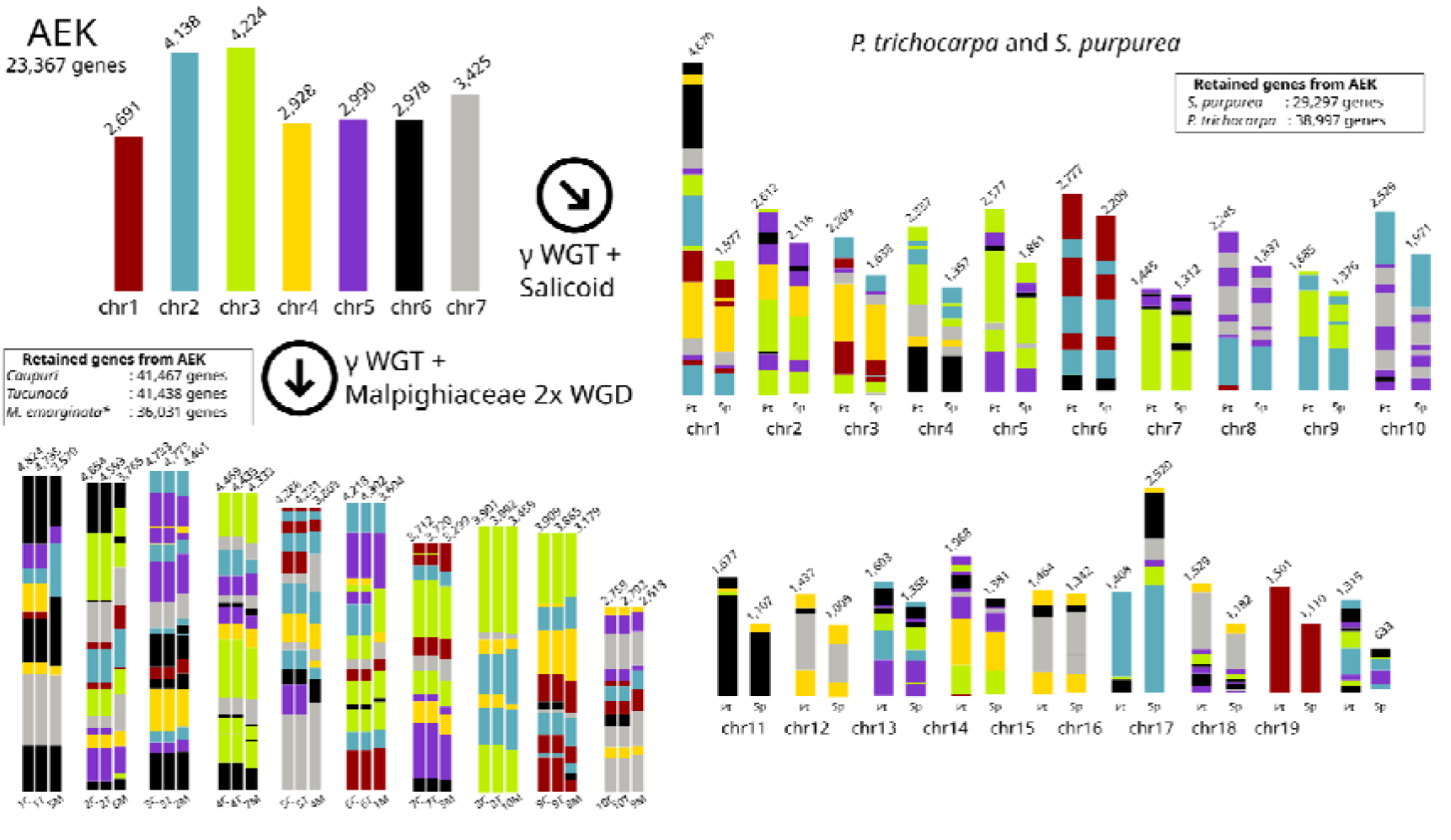
Reconstruction of the ancestral eudicot karyotype (AEK) and polyploidization history of *Banisteriopsis caapi* and related species using WGDI. The AEK consisted of seven protochromosomes (upper-left panel) that harbored 23,367 ancestral genes. Gene retention patterns following subsequent polyploidization events are illustrated for *B. caapi* morphotypes (caupuri and tucunacá) and *Malpighia emarginata* (bottom left), as well as for *Populus trichocarpa* and *Salix purpurea* (right panel). The analyses support the occurrence of the γ whole-genome triplication (WGD), followed by lineage-specific WGDs: the Salicoid event in *Populus* and *Salix*, and at least two WGDs within Malpighiaceae. Numbers above bars indicate the number of genes retained from the AEK in each species/chromosome, shown below the bars. The chromosome bars are not to scale and are grouped by pattern similarity.

### 3.5. Transposable element-associated insertions and deletions dominate the structural-variation (SVs) landscape

Reciprocal whole-genome comparisons between the caupuri and tucunacá assemblies identified 14,361 structural variants (SVs ≥ 50 bp), collectively spanning approximately 20% of each assembly when translocation spans were excluded from genome-coverage estimates (Figure 4A,B; Table S8). Transposable element-associated insertions and deletions represented the most abundant SV classes. Mobile insertions accounted for 6,089 events and approximately 107 Mb, whereas mobile deletions accounted for 5,760 events and approximately 94 Mb. Novel insertions and deletions, defined here as events without overlap with annotated TEs, were less frequent, comprising 299 insertion events and 335 deletion events in the caupuri-oriented comparison, and 329 insertion events and 303 deletion events in the tucunacá-oriented comparison (Table S8).

**Figure 4.**
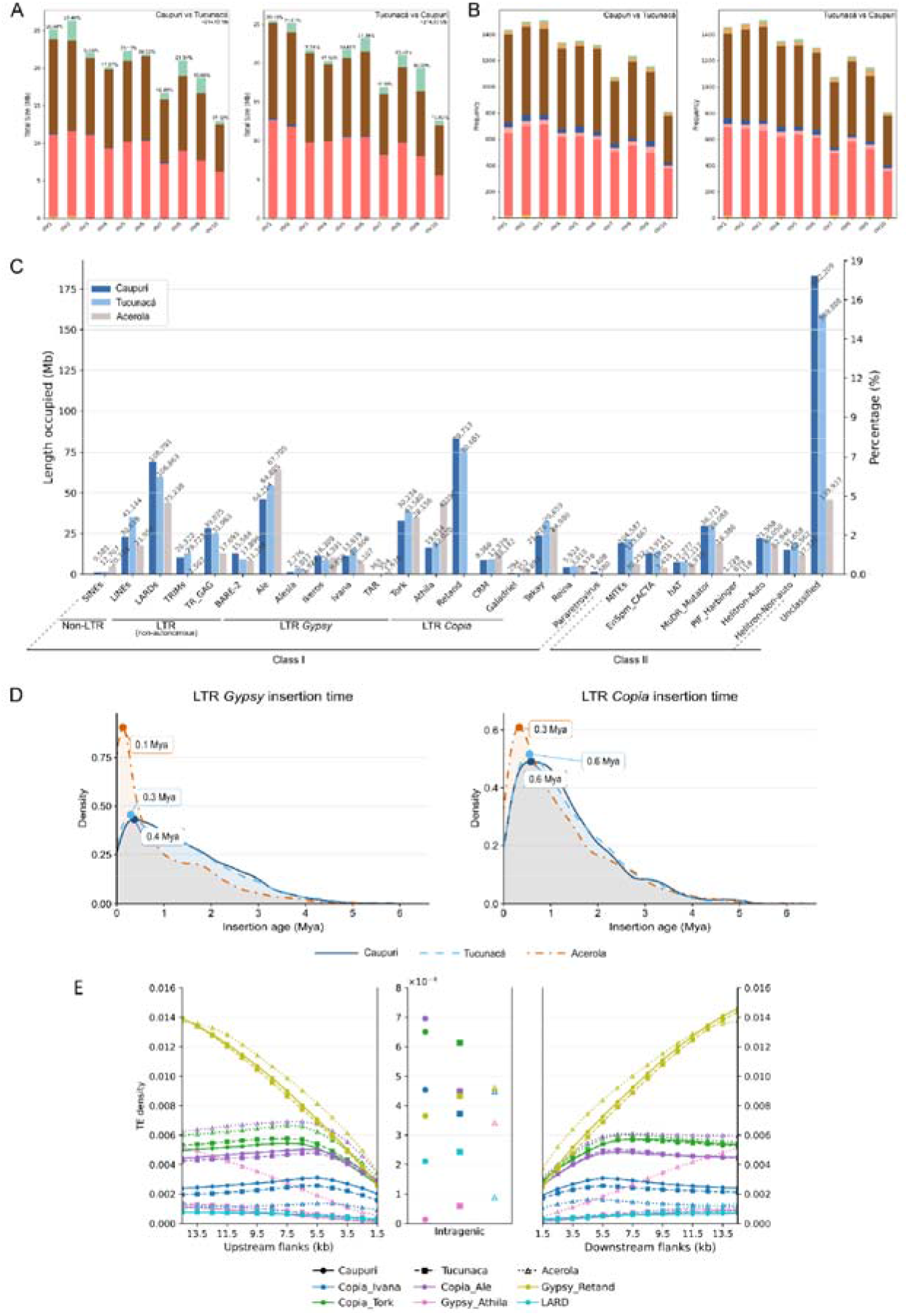
Structural variation, transposable element composition, LTR insertion history, and gene-centered TE density in *Banisteriopsis caapi* morphotypes. (A–B) Total cumulative size (A) and frequency (B) of structural variants (SVs) identified in reciprocal whole-genome comparisons between caupuri and tucunacá. Bars show SV frequencies for insertions, deletions, inversions, duplications, and translocations. (C) Comparative transposable element composition of caupuri, tucunacá, and acerola (*Malpighia emarginata*). The left y-axis indicates the cumulative genomic length occupied by each TE category, whereas the right y-axis indicates the corresponding percentage of genome occupancy. Numbers above bars indicate annotated element counts. Colors follow the morphotype legend (top left). (D) Kernel-density distributions of intact LTR/*Gypsy* and LTR/*Copia* insertion ages in caupuri, tucunacá, and *M. emarginata*. Colors and line types identify the three genomes, and labeled points indicate the dominant density maximum of each distribution. Insertion ages are shown in millions of years ago and were estimated using a neutral mutation rate of 1.1 × 10 substitutions per site per year. Complete genome-specific repeat-divergence landscapes and LTR insertion-time distributions are presented in Figure S8. (E) Average density profiles of selected LTR-related categories and LARD elements across upstream flanking regions, annotated gene bodies, and downstream flanking regions in caupuri, tucunacá, and *M. emarginata*.

Other SV classes were recovered at lower frequencies. Across the reciprocal comparisons, duplications comprised 371 events in the caupuri-oriented comparison and 318 in the tucunacá-oriented comparison, whereas inversions comprised 118 and 126 events, respectively. Contractions accounted for 101 events in the caupuri-oriented comparison and 117 in the tucunacá-oriented comparison. Translocations were also detected in both directions of comparison, with 451 events in the caupuri-oriented comparison and 469 in the tucunacá-oriented comparison. Because cumulative translocation spans can be inflated by large-scale alignment breaks, these spans were reported separately from the total occupied genomic fraction (Table S8).

SVs were distributed across all chromosomes in both assemblies (Figure S7; Table S8). The highest total SV counts occurred on chromosomes 3, 2, and 1, whereas the lowest count occurred on chromosome 10. Because these counts were not normalized by chromosome length, chromosome-wise differences were interpreted descriptively. Chromosome-wise occupied genomic fractions ranged from 16.41% to 21.54% in caupuri and from 16.11% to 21.34% in tucunacá, excluding translocation spans from the total coverage estimates.

### 3.6. Repeat landscapes are highly similar in *B. caapi* but compositionally distinct from

### Malpighia emarginata

The three Malpighiaceae assemblies showed similar total repetitive fractions, accounting for 77.96% of the caupuri assembly, 78.27% of the tucunacá assembly, and 77.87% of the *M. emarginata* assembly (Figure 4C; Table 2; Table S9). In both *B. caapi* morphotypes, Class I retrotransposons represented the largest repeat fraction, occupying 55.47% of the caupuri assembly and 57.24% of the tucunacá assembly. LTR/*Gypsy* elements accounted for 28.39% and 28.33% of the two assemblies, respectively, whereas LTR/*Copia* elements occupied 13.30% of caupuri and 15.07% of tucunacá. Non-autonomous LTR derivatives represented 10.43% of the caupuri assembly and 9.52% of the tucunacá assembly, with LARDs forming the largest fraction within this group, corresponding to 5.99% and 5.30% of the two assemblies, respectively. Non-LTR retrotransposons occupied 2.09% of the caupuri assembly and 3.27% of the tucunacá assembly, with the difference mainly attributable to LINE content. Class II DNA transposons represented 7.69% of the caupuri assembly and 8.28% of the tucunacá assembly, including TIR elements, MITEs, and Helitrons. *M. emarginata* showed a comparable total repeat fraction but differed in class composition, with lower non-autonomous LTR content (6.82%), lower LARD content (4.21%), and a larger Class II fraction (18.64%). The unclassified repeat fraction was 15.82% in caupuri, 14.09% in tucunacá, and 3.42% in *M. emarginata*, indicating a substantial component of repeat sequences in *B. caapi* that could not be assigned to known categories (Figure 4C; Table 2; Table S9).

Among LTR lineages and related categories, the two *B. caapi* morphotypes showed broadly similar profiles, with quantitative differences in specific components (Figure 4C). Tucunacá showed higher genome occupancy of *Copia*/Ale (4.82% versus 4.01%), *Copia*/Ivana (1.41% versus 0.97%), and *Copia*/Tork (3.41% versus 2.85%), whereas caupuri showed higher LARD content (5.99% versus 5.30%) and slightly higher *Gypsy*/Retand representation (7.21% versus 6.82%). Relative to both *B. caapi* morphotypes, *M. emarginata* showed higher contributions from *Copia*/Ale (6.14%), *Gypsy*/Athila (4.89%), and *Gypsy*/Retand (8.34%), and lower *Copia*/Ivana content (0.61%).

### 3.7. Repeat-divergence landscapes, LTR insertion times, and gene-centered TE density

Repeat landscapes based on Kimura divergence showed similar profiles for the two *B. caapi* morphotypes, with the largest contribution deriving from LTR/*Gypsy* elements, followed by LTR/*Copia* elements (Figure 4D and Figure S8). In both morphotypes, repeat annotations were concentrated mainly in low-to-intermediate Kimura-divergence classes and decreased toward higher divergence values. *M. emarginata* showed a distinct repeat-divergence profile, with a larger Class II contribution and a lower unclassified repeat fraction than the two *B. caapi* morphotypes (Figure S8).

Estimated insertion-time distributions of intact LTR retrotransposons showed profiles that closely corresponded between the two *B. caapi* ethnovarieties (Figure 4D and Figure S8). For LTR/*Gypsy* elements, caupuri and tucunacá showed mean insertion times of 1.38 and 1.34 Mya, median insertion times of 1.17 and 1.14 Mya, and dominant density maxima at approximately 0.4 and 0.3 Mya, respectively. In *M. emarginata*, the LTR/*Gypsy* distribution was shifted toward younger ages, with a mean of 0.90 Mya, a median of 0.52 Mya, and a dominant density maximum at approximately 0.1 Mya. LTR/*Copia* elements showed more similar distributions across the three genomes. Caupuri and tucunacá had mean insertion times of 1.31 and 1.30 Mya, median insertion times of 1.08 and 1.07 Mya, and dominant density maxima at approximately 0.6 Mya in both genomes. In *M. emarginata*, LTR/*Copia* elements showed a mean insertion time of 1.17 Mya, a median of 0.88 Mya, and a dominant density maximum at approximately 0.3 Mya. Thus, the two *B. caapi* genomes showed nearly overlapping *Gypsy* and *Copia* insertion-time distributions, whereas *M. emarginata* displayed a stronger young insertion component, particularly for *Gypsy* elements.

Genome-wide TE-density profiles around genes were generated for selected LTR-related categories and LARD elements in caupuri, tucunacá, and *M. emarginata* (Figure 4E). For *Copia*/Ale, *Copia*/Ivana, *Copia*/Tork, *Gypsy*/Athila, *Gypsy*/Retand, and LARD elements, TE densities were lower within gene bodies than in upstream and downstream flanking regions. Flanking TE-density values generally increased with increasing distance from gene bodies. The same categories were also examined in chromosome-resolved profiles (Figure S9). Across chromosome-resolved profiles, *Gypsy*/Retand generally showed the highest TE-density values among the selected categories. In the two *B. caapi* morphotypes, *Copia*/Ale and *Copia*/Tork generally occupied intermediate profiles, whereas *Copia*/Ivana, *Gypsy*/Athila, and LARD were lower in most chromosome-level comparisons. *Malpighia emarginata* showed a similar gene-centered profile structure, although chromosome-level profiles differed in the relative magnitude of *Copia*/Ale, *Copia*/Tork, and *Gypsy*/Athila densities.

### 3.8. Gene-family evolution in chromosome-anchored *B. caapi* gene sets

OrthoFinder assigned most chromosome-anchored genes from the two *B. caapi* morphotypes to shared orthogroups. A total of 25,004 orthogroups were shared between the two sampled genomes, while assembly-restricted orthogroups were substantially less prevalent. A total of 350 orthogroups containing 1,012 genes were represented only in the caupuri assembly, while 316 orthogroups containing 791 genes were represented only in the tucunacá assembly across the complete comparative dataset (Table S12). The most recurrent representative annotations across distinct assembly-restricted orthogroups were aminotransferase-like plant mobile-domain proteins, serine/threonine-protein phosphatase 7 homologs, and XRCC3-like DNA-repair proteins in caupuri, and aminotransferase-like plant mobile-domain proteins, serine/threonine-protein phosphatase 7 homologs, and ADP-ribosyl cyclase/cyclic ADP-ribose hydrolases in tucunacá (Table S12). This pattern indicates that the restricted orthogroups are concentrated in a limited number of repeatedly represented protein classes rather than being broadly distributed across unrelated functions.

For CAFE5 analysis, families present in only one species and families with ≥100 copies in at least one species were removed, resulting in 29,850 gene families submitted to the final analysis. The calibrated CAFE5 model estimated λ = 0.002704 and ε = 0.017197. CAFE5 reconstructed 286 gene-family expansions and 246 contractions in caupuri, and 381 expansions and 203 contractions in tucunacá (Figure 5A; Table S14). Branch-specific probability testing identified a smaller high-confidence subset, comprising 29 significant expansions and 25 significant contractions in caupuri, and 40 significant expansions and 28 significant contractions in tucunacá. Thus, tucunacá retained a stronger terminal expansion signal than caupuri.

**Figure 5.**
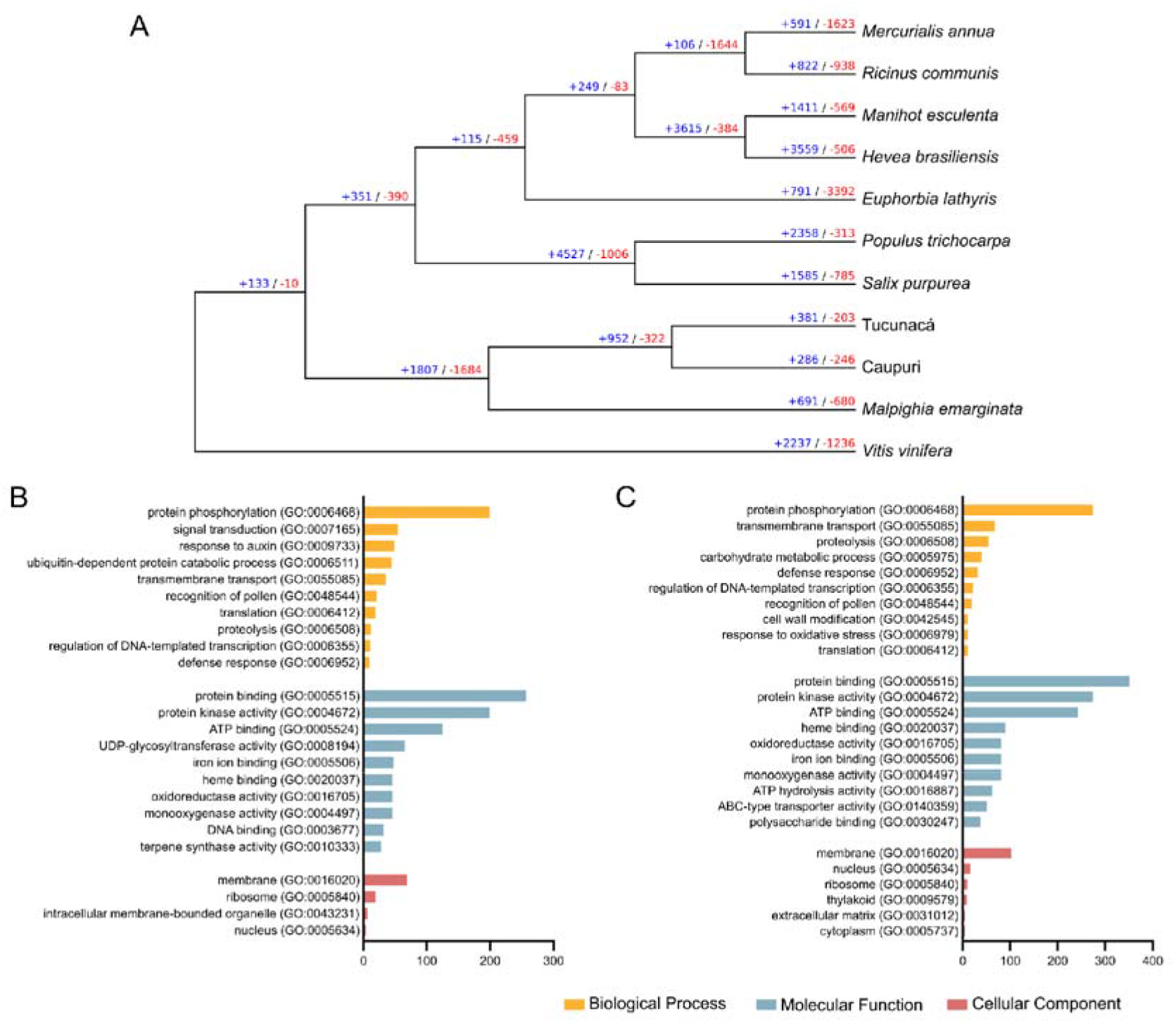
Gene family evolution and functional enrichment in *Banisteriopsis caapi* morphotypes. (A) CAFE5-reconstructed gene-family increases and decreases across the calibrated species tree. Values indicate reconstructed copy-number changes in each branch. Blue and red numbers indicate gene family expansions and contractions, respectively. (B–C) Gene Ontology enrichment analysis of genes belonging to expanded orthogroups in the terminal caupuri (B) and tucunacá (C) branches.

GO enrichment analysis of genes from expanded orthogroups revealed a broadly shared functional background, with partially distinct emphases (Figure 5B and C; Table S14). Expanded families in both morphotypes were associated with protein phosphorylation, transmembrane transport, proteolysis, transcriptional regulation, pollen recognition, and defense response, indicating recurrent remodeling of regulatory, transport, and environmental-response functions (Figure 5B and C). Within this shared profile, the caupuri enrichment highlighted signal transduction, auxin response, ubiquitin-dependent protein turnover, UDP-glycosyltransferase activity, and terpene synthase activity (Figure 5B), whereas the tucunacá profile included carbohydrate metabolism, cell wall modification, oxidative-stress response, ABC-type transporter activity, and polysaccharide binding (Figure 5C). These independently inferred enrichment profiles suggest that gene-family expansion affected a common regulatory and metabolic framework in both genomes, while emphasizing different subsets of hormone-, specialized-metabolism-, stress-, and cell-wall-associated functions.

Pairwise OrthoVenn3 analysis of the chromosome-anchored protein sets recovered 30,773 clusters, of which 30,181 (98.1%) were shared between caupuri and tucunacá (Table S15). Shared clusters contained 33,630 caupuri genes and 33,573 tucunacá genes, whereas 259 caupuri-only clusters contained 723 genes and 333 tucunacá-only clusters contained 893 genes. The predominance of shared clusters independently supports the conserved gene-family background identified by OrthoFinder. To connect cluster exclusivity with phylogenetically informed copy-number changes, genes from the exclusive clusters were mapped to their OrthoFinder orthogroups and corresponding CAFE5 results. Twenty-seven caupuri-only and 46 tucunacá-only clusters overlapped prioritized CAFE5 candidate orthogroups, and the direction of the CAFE5 change was concordant with cluster exclusivity for 23 and 42 clusters, respectively. After excluding clusters dominated by TE-related, organellar, or uninformative annotations, 20 caupuri and 40 tucunacá clusters were retained as integrated candidates; four caupuri and nine tucunacá clusters also overlapped orthogroups with branch-specific significant CAFE5 changes. These candidates included MADS-box transcription factors, receptor-like and wall-associated kinases, ABC transporters, UDP-glycosyltransferases, bHLH-related transcription factors, and genes associated with defense and mechanical or cell-wall perception. OrthoVenn3 clusters detected only in one genome mapped to OrthoFinder orthogroups that also contained shared OrthoVenn3 clusters. In these cases, cluster exclusivity represents a genome-biased subdivision or copy-number difference within a gene family present in both assemblies, rather than a gene family restricted entirely to one genome. The highest-confidence loci emerging from the integrated CAFE5, OrthoVenn3, tandem-array, and functional-prioritization analyses are summarized in Table 3.

**Table 3.** Prioritized candidate genes and local gene clusters associated with stem-architecture divergence between caupuri and tucunacá.

| Candidate locus or gene group | Representative genes/loci | Evidence | Candidate relevance |
| --- | --- | --- | --- |
| MADS-box developmental TFs in Caupurí and Tucunacá | CaupuriG00000024047/24050/24057;<br>TucunacaG00000062572/62580/62581/62586 | Expanded candidate OGs; developmental and shoot/stem categories | Candidate regulators of stem and shoot developmental programs |
| B3/VRN1-like TF tandem array in Caupurí | CaupuriG00000052282-CaupuriG00000052286 | Large tandem array; expanded candidate OG; transcriptional regulation | Local expansion of developmental regulatory genes in Caupurí |
| Wall-associated receptor kinase loci | CaupuriG00000012488/12489;<br>TucunacaG00000007999/08000;<br>TucunacaG00000041778/41781/41784/41795 | Specific or expanded candidate clusters; WAK/Pkinase domains | Candidate cell-wall perception and signaling genes |
| Mechanosensitive ion-channel tandem pair in Tucunacá | TucunacaG00000016137/16138 | Expanded candidate OG; tandem topology; developmental categories | Candidate genes for mechanical perception and cell expansion |
| Cell-wall remodeling and biosynthesis genes | CaupuriG00000044833/44834;<br>CaupuriG00000010276/10278/10279;<br>TucunacaG00000031936/33052/33066 | Cell wall-related candidate OGs; tandem or expanded loci | Candidates for wall biosynthesis, pectin modification, and stem tissue organization |
| Transport-associated candidate clusters | CaupuriG00000067204-CaupuriG00000067218;<br>TucunacaG00000023315/23320/23323/23326 | Expanded candidate OGs; transport, hormone, and developmental categories | Potential regulators of metabolite, hormone, or wall-precursor transport during stem differentiation |
| High-copy tandem arrays with wall/defense-related annotations | CaupuriG00000012889-CaupuriG00000012920;<br>TucunacaG00000062430-TucunacaG00000062434 | Largest Caupurí array, 33 genes; large Tucunacá array, 15 genes | Focal local expansions affecting defense-, wall-, and cell-wall-modification-related functions |

### 3.9. Tandem-duplication landscapes in the sampled *B. caapi* genomes

Tandem-duplicated genes were detected in both sampled *B. caapi* genomes and examined by integrating tandem-array structure, Ks estimates for tandem gene pairs, and enrichment analyses of tandem-classified genes. The curated tandem-array set comprised 228 arrays in caupuri and 238 in tucunacá (Table S16; Figure 6A). These tandem pairs were distributed unevenly across all chromosomes in both ethnovarieties. In caupuri, tandem pairs were most frequent on chromosomes 2, 1, and 7, with 31, 30, and 27 arrays, respectively. Tucunacá contained the largest numbers on chromosomes 1, 6, and 2, with 41, 31, and 28 arrays. Most arrays were small, especially two- and three-gene arrays, indicating that tandem duplications in these assemblies were predominantly organized as short local clusters. Large arrays were less frequent but included a 33-gene endochitinase-related array on caupuri chromosome 2 and a 15-gene array containing pectin methylesterase inhibitor- and receptor-like kinase-related annotations in tucunacá. Thus, although the overall distribution was dominated by small arrays, both genomes retained localized high-copy tandem clusters with annotations related to cell wall modification, signaling, and defense-associated enzymatic activity.

**Figure 6.**
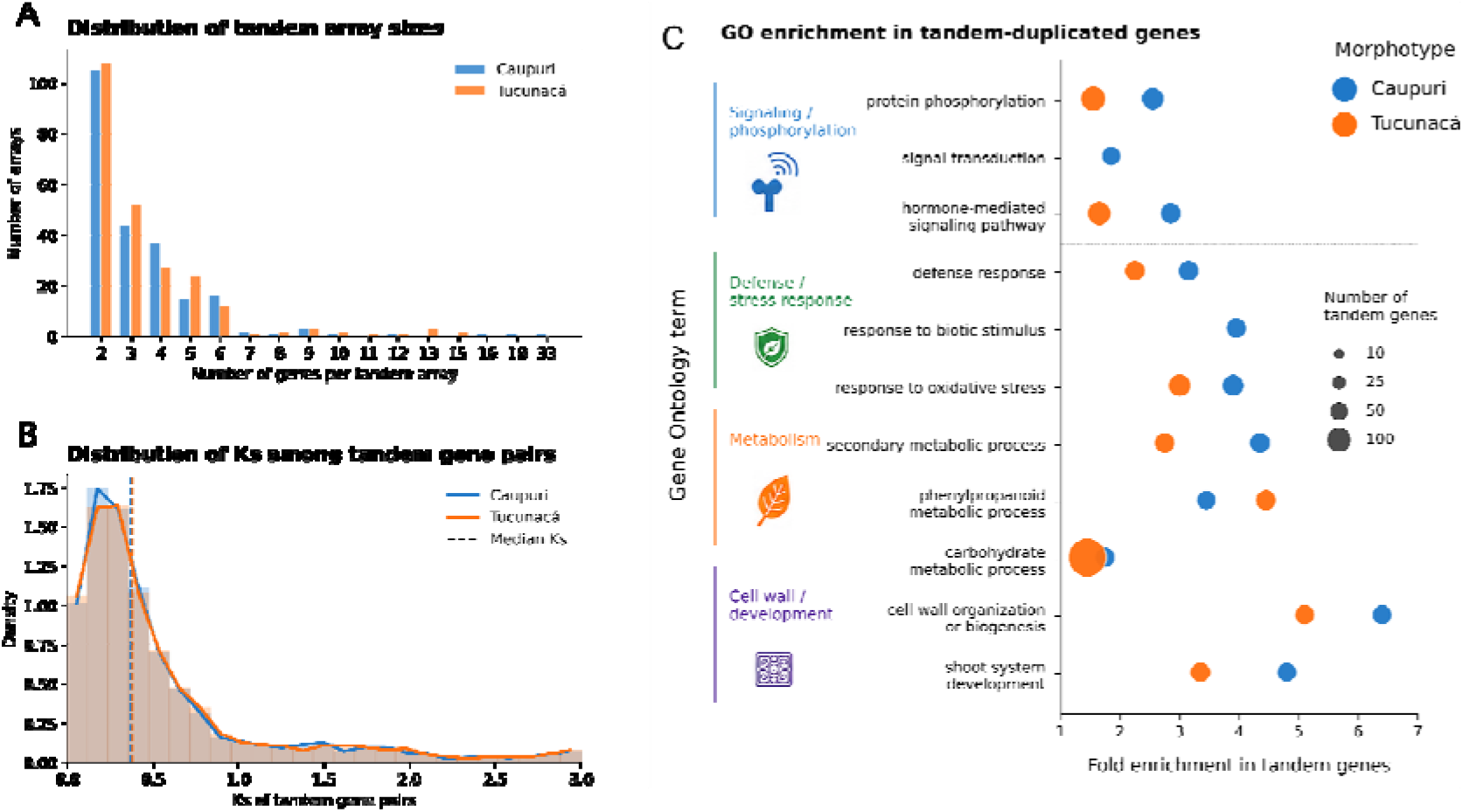
Tandem-duplication architecture, Ks distribution, and functional enrichment of tandem-duplicated genes in the caupuri and tucunacá morphotypes of *Banisteriopsis caapi*. (A) Distribution of tandem-array sizes in the two morphotypes. Bars represent the number of tandem arrays observed for each array size, without grouping of large arrays. Both morphotypes were dominated by small arrays, but large local clusters were also detected, including a maximum of 33 genes in caupuri and 15 genes in tucunacá. (B) Ks distribution of tandem gene pairs with valid estimates. Histograms and density curves are shown for visualization at Ks ≤ 3, and dashed vertical lines represent median Ks values for each morphotype. The nearly overlapping distributions indicate similar global age structures of tandem duplicates in the two genomes. (C) Representative GO terms enriched among tandem-duplicated genes. The x-axis shows fold enrichment relative to the annotated genomic background, point size represents the number of tandem genes assigned to each term, and color denotes morphotype. Displayed terms summarize significant categories related to signaling and phosphorylation, defense and stress response, secondary and carbohydrate metabolism, cell wall organization, and shoot-system development. Complete GO enrichment results and complementary COG and EC enrichment analyses are listed in Table S17.

Ks estimates were then evaluated for adjacent gene pairs reconstructed within tandem arrays. Valid Ks estimates were obtained for 3,046 pairs in caupuri, with median Ks = 0.375 and mean Ks = 0.914, and for 3,087 pairs in tucunacá, with median Ks = 0.382 and mean Ks = 1.045 (Figure 6B). The two distributions were nearly overlapping, and most pairs were concentrated at low Ks values. In caupuri, 62.2% of valid tandem pairs showed Ks < 0.5 and 79.0% showed Ks < 1.0. In tucunacá, 61.2% showed Ks < 0.5 and 79.0% showed Ks < 1.0. Overall, the two genomes showed similar tandem-array size distributions and tandem-pair Ks profiles, with substantial contributions from low-Ks pairs consistent with relatively recent local duplications.

GO enrichment of tandem-duplicated genes revealed significant overrepresentation of terms associated with signaling, stress response, metabolism, cell wall organization, and development (Figure 6C; Table S17). Enriched terms included protein phosphorylation, signal transduction, hormone-mediated signaling pathway, defense response, response to biotic stimulus, response to oxidative stress, secondary metabolic process, phenylpropanoid metabolic process, carbohydrate metabolic process, cell wall organization or biogenesis, and shoot system development. These terms indicate that tandem-duplicated genes are not randomly distributed across functional annotation classes but are enriched in categories relevant to regulatory responses, environmental interaction, specialized metabolism, cell wall dynamics, and shoot-system development. Complementary COG and EC enrichments were consistent with the GO-based pattern, further supporting an enrichment of tandem-duplicated genes in regulatory, metabolic, defense-related, and cell wall-associated annotation categories (Table S17). Pairwise comparisons between caupuri and tucunacá using chromosome-anchored annotated gene sets and tandem-duplicated gene subsets further indicated that these enrichment profiles were not solely explained by broad differences in genome-wide annotation composition. The recurrent enrichment of signaling, defense, metabolic, cell-wall-related, and developmental functions was therefore associated specifically with the tandem-duplicated gene sets in both sampled genomes.

Together, the array size, Ks, and enrichment results indicate that caupuri and tucunacá share a broadly similar tandem-duplication landscape, characterized by mostly small arrays and nearly overlapping low-Ks distributions. Against this similar overall background, both genomes retained localized high-copy arrays, and their tandem-duplicated gene sets showed recurrent enrichment in functional categories associated with signaling, stress response, metabolism, cell wall biology, and development.

### 3.10. LARD–gene associations and functional enrichment

To integrate the repeat-landscape and local-duplication results, we examined the positional, functional, and duplication-related associations between LARD elements and annotated genes. LARD–gene associations were classified as gene_inside_LARD, LARD_inside_gene, partial_overlap, or flank-only according to the positional relationship between LARD annotations and gene coordinates (Figure 7; Table S11). Before gene-level collapsing, complete gene containment within LARD annotations was detected for 10 genes in caupuri, four genes in tucunacá, and one gene in *M. emarginata*. Associations in which a LARD interval was entirely contained within an annotated gene involved 1,005 genes in caupuri, 859 genes in tucunacá, and 16 genes in *M. emarginata*. Partial overlaps involved 373 genes in caupuri and 249 genes in tucunacá and were not recovered in *M. emarginata* under the same criteria. Flank-only associations within 5 kb of annotated genes involved 15,391 genes in caupuri, 15,558 genes in tucunacá, and 77 genes in *M. emarginata*.

**Figure 7.**
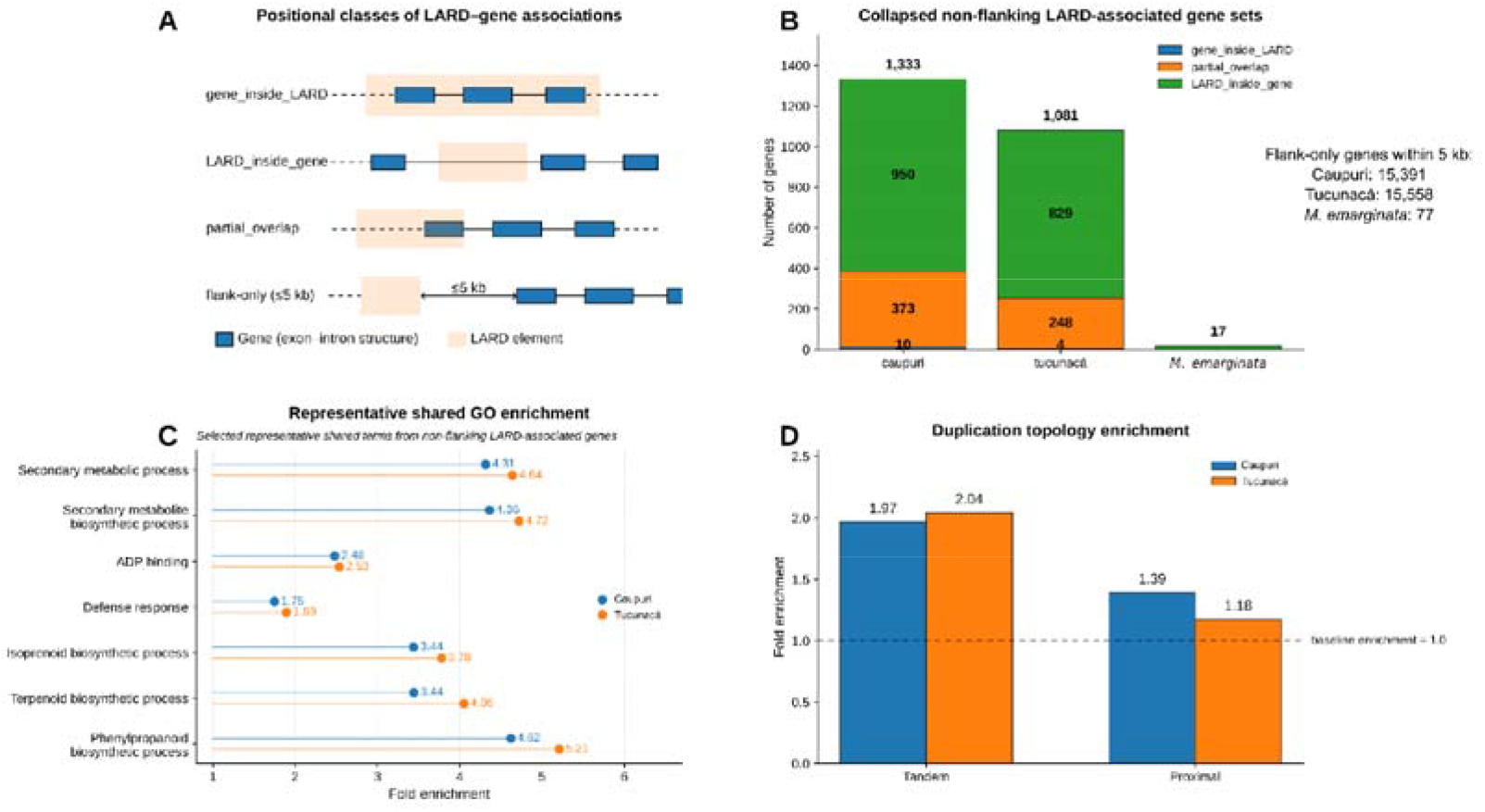
Positional, functional, and duplication-related patterns of genes associated with large retrotransposon derivative elements in *Banisteriopsis caapi.* (A) Schematic representation of the four positional relationships evaluated between annotated genes and large retrotransposon derivative (LARD) elements: gene_inside_LARD, in which the complete gene interval is contained within a LARD annotation; LARD_inside_gene, in which the complete LARD interval is contained within the gene span; partial_overlap, in which the gene and LARD intervals overlap but neither is completely contained within the other; and flank-only, defined as a non-overlapping LARD located within 5 kb upstream or downstream of a gene. (B) Numbers of genes assigned to the mutually exclusive non-flanking LARD-association classes after each gene was collapsed to its strongest positional relationship. Flank-only associations are reported separately because their substantially larger numbers would obscure the non-flanking categories. *Malpighia emarginata* was not included in the functional-enrichment analyses because only 17 non-flanking LARD-associated genes were recovered and an equivalent integrated functional-annotation background was unavailable. (C) Representative Gene Ontology terms significantly enriched among non-flanking LARD-associated genes in caupuri and tucunacá. Points and numeric labels indicate fold enrichment relative to the corresponding chromosome-level annotated gene background. Only representative terms shared between the two morphotypes are shown; complete enrichment results are provided in Table S11. (D) Enrichment of tandem- and proximal-duplication topologies among non-flanking LARD-associated genes. Bars indicate fold enrichment relative to the corresponding genome-wide duplication-topology background, and the dashed horizontal line represents the null expectation of fold enrichment = 1. GO, COG, EC, and duplication-topology profiles did not differ significantly between caupuri and tucunacá after Benjamini–Hochberg false-discovery-rate correction.

After collapsing each gene to its strongest non-flanking LARD association, the resulting dataset comprised 1,333 genes in caupuri, 1,081 genes in tucunacá, and 17 genes in *M. emarginata* (Figure 7; Table S11). In both *B. caapi* morphotypes, this set was dominated by LARD_inside_gene and partial_overlap relationships. Functional enrichment analyses were performed independently for caupuri and tucunacá using all genes carrying the corresponding annotation type on the chromosome-level assemblies as background. *M. emarginata* was not included in the enrichment analysis because only 17 non-flanking genes were recovered and an equivalent integrated functional-annotation background was unavailable.

GO enrichment identified 78 significant terms in caupuri and 65 in tucunacá. The two morphotypes showed broadly concordant enrichment profiles, including secondary-metabolite biosynthesis, defense response, ADP binding, signal transduction, isoprenoid and terpenoid metabolism, phenylpropanoid metabolism, xenobiotic transport and detoxification, and oxidoreductase activity (Figure 7; Table S11). COG analyses similarly recovered enrichment for secondary-metabolite biosynthesis, transport, and catabolism; defense mechanisms; and lipid transport and metabolism in both morphotypes. At the gene-description level, recurrent annotations among the non-flanking LARD-associated genes included receptor-like kinases, leucine-rich repeat proteins, F-box proteins, ADP-ribosyl cyclase/cyclic ADP-ribose hydrolases, and resistance-like proteins (Table S11). Non-flanking LARD-associated genes were also enriched for tandem duplication, with fold-enrichment values of 1.97 in caupuri and 2.04 in tucunacá, and for proximal duplication, with corresponding values of 1.39 and 1.18 (Figure 7; Table S11). No significant differences between caupuri and tucunacá were recovered for GO, COG, EC, or duplication-topology profiles after multiple-testing correction. Together, these results show that non-flanking LARD-associated genes are nonuniformly represented across functional and duplication categories, with enrichment for locally duplicated genes and functions related to defense, environmental signaling, transport, and specialized metabolism. The absence of significant differences between the morphotypes indicates that this pattern represents a shared feature of the *B. caapi* genomic architecture rather than a morphotype-specific enrichment, although individual LARD-associated loci may still contribute to gene-level functional divergence.

## 4. Discussion

This study generated chromosome-scale nuclear genomic resources for the caupuri and tucunacá morphotypes of *B. caapi*, providing a framework to examine genome evolution, structural variation, repeat dynamics, and candidate genomic features associated with morphotype-level differentiation in this culturally important Amazonian liana. Within this framework, the two sampled genomes appear to have evolved within a largely shared paleopolyploid genomic background, despite the low chromosome number reported for the species (2n = 20) [85]. Low chromosome numbers are not necessarily incompatible with ancestral genome duplication, as descending dysploidy has been proposed for other Malpighiales lineages [86]. Synteny, Ks distributions, WGD analyses, and ancestral eudicot karyotype reconstruction recovered older duplication components in both morphotypes, with comparable signals also detected in *M. emarginata* [11, 82]. These results suggest that caupuri, tucunacá and *M. emarginata* retain signatures of a shared paleopolyploid history. However, broader sampling within Malpighiaceae will be required to determine the taxonomic distribution and phylogenetic placement of these duplication signals [82].

Within this shared genomic background, structural variation and repetitive elements reveal a second layer of differentiation between the sampled genomes. Caupuri and tucunacá have similar assembly sizes, total repeat fractions, and gene-centered TE-density profiles, and nearly overlapping LTR insertion-time distributions, indicating that they are not distinguished by broad differences in chromosome number, genome size, or large-scale TE organization. The dominant young components recovered for both *Gypsy* and *Copia* elements, concentrated at approximately 0.3–0.6 Mya in the two *B. caapi* genomes (Figure 4), provide evidence of a shared recent-to-intermediate history of LTR activity rather than strongly morphotype-specific retrotransposon bursts. By comparison, *M. emarginata* showed a more pronounced young component, particularly for *Gypsy* elements, indicating lineage-level variation in the temporal dynamics of LTR accumulation. Caupuri and tucunacá nevertheless differed quantitatively in specific repeat compartments, including LTR-related lineages, LINEs, Class II elements, LARDs, and the unclassified repeat fraction. Most structural variants corresponded to mobile element-associated insertions and deletions, suggesting that TE-related turnover contributed substantially to local genomic differentiation between the two assemblies. Although the insertion-time distributions do not associate individual structural variants with particular dated LTR elements, the presence of young *Gypsy* and *Copia* components is consistent with comparatively recent repeat turnover capable of contributing to localized structural remodeling. Such variation is consistent with the role of TE as generators of structural polymorphism and regulatory variation in plant genomes [13, 15]. Thus, repetitive elements and SVs appear to differentiate the sampled genomes mainly through localized and quantitative remodeling rather than through extensive genome-wide reorganization.

Gene-family evolution provides an additional layer for interpreting differences between the sampled caupuri and tucunacá genome assemblies. Most chromosome-anchored genes were assigned to shared orthogroups, whereas assembly-restricted orthogroups represented a comparatively small fraction of the total repertoire. This pattern indicates that the sampled genomes differ mainly through copy-number shifts and differential remodeling within shared gene families, rather than through the emergence of extensive morphotype-exclusive repertoires. Such a pattern is consistent with plant-genome studies showing that copy-number variation and other structural variants can contribute to phenotypic diversification and environmental adaptation through dosage changes, presence–absence variation, and regulatory alterations affecting particular loci or functionally enriched subsets of genes [87, 88]. Consistent with this interpretation, CAFE5 recovered asymmetric remodeling in the terminal branches, with a stronger terminal expansion signal in tucunacá. The OrthoVenn3 analysis provided complementary support: most protein clusters were shared between the two assemblies, while a smaller set of exclusive clusters overlapped expanded or contracted CAFE5 orthogroups in a direction consistent with the genome in which they were detected. Because several of these exclusive clusters belong to broader OrthoFinder orthogroups represented in both assemblies, they more likely reflect localized differentiation within shared gene families than complete gain or loss of entire families. The strongest integrated candidates included MADS-box transcription factors, receptor-like and wall-associated kinases, transporters, glycosyltransferases, and genes associated with defense, mechanical perception, and cell wall biology. Together, these analyses identify a restricted set of candidate gene families and protein clusters embedded within a predominantly conserved gene-family background.

The LARD–gene association analyses revealed a shared relationship between LARD-containing gene regions and local gene duplication in the two *B. caapi* genomes. LARDs were originally characterized as abundant, dispersed, and nonautonomous LTR-retrotransposon derivatives lacking the protein-coding capacity of autonomous elements [89]. In the caupuri and tucunacá assemblies, non-flanking LARD-associated genes were significantly overrepresented among tandem duplicates, with approximately two-fold enrichment in both genomes, and showed additional, weaker enrichment among proximal duplicates. The LARD-associated and tandem-duplicated gene sets also showed overlapping broad functional categories, particularly defense, signaling, secondary metabolism, and phenylpropanoid metabolism. This convergence does not demonstrate that LARD insertions caused local gene duplication or altered the regulation of the associated genes, but it identifies a recurrent relationship between LARD-containing regions and locally duplicated functional modules. Such a relationship is compatible with evidence that LTR-retrotransposon-mediated gene duplication can be followed by tandem amplification of defense-related genes [90] and that TEs located within or near genes can become associated with stress-responsive genomic networks [91]. Because the functional and duplication-topology profiles did not differ significantly between caupuri and tucunacá, this pattern is best interpreted as a shared feature of *B. caapi* genome architecture rather than as an ethnovariety-specific process.

Tandem duplication added a more recent and local layer to this pattern. Caupuri and tucunacá showed highly similar global tandem architecture, with mostly small arrays and nearly overlapping Ks distributions. Within this broadly comparable background, each genome retained a limited number of localized high-copy arrays, while tandem-duplicated genes were enriched for functions related to signaling, defense, stress response, metabolism, cell wall biology, and development. The predominance of low-Ks tandem pairs is consistent with local expansions that are comparatively recent relative to the older paleopolyploid background, in agreement with the view that tandem duplication provides a recurrent, small-scale source of gene-family remodeling after genome-wide duplication events [16, 92]. Arrays detected only in one sampled assembly may reflect accession-specific local expansions, but broader population sampling will be required to determine whether they are consistently associated with ethnovariety differentiation. This pattern is consistent with broader plant-genome studies showing that tandemly arrayed genes are enriched for membrane-associated and stress-related functions, and that lineage-specific tandem expansions often contribute to environmental-response and biotic-stress gene repertoires [16, 93–95]. Thus, tandem duplication may represent a post-polyploid, focal amplification process affecting candidate developmental, signaling, transport, and wall-related modules in both sampled genomes.

The indirect estimate of approximately 1.13 Ma for the caupuri–tucunacá node provides additional temporal context for this localized genomic remodeling. Median intact-LTR insertion ages in the two genomes were close to this timescale, whereas the dominant *Gypsy* and *Copia* density maxima at approximately 0.3–0.6 Ma indicate substantial LTR activity during a more recent period. Tandem-pair Ks distributions also showed a strong low-Ks component, indicating local duplications that are comparatively recent relative to the ancient WGD background. Together with the enrichment of tandem duplicates among LARD-associated genes, these results are compatible with continued repeat turnover and local gene-family remodeling during the recent evolutionary history of the sampled accessions. However, individual tandem arrays and LARD insertions cannot be assigned as occurring before or after their separation from the available data.

Taken together, the conservation of genome size, chromosome number, chromosome-scale collinearity, paleopolyploid background, and shared gene repertoire, combined with differences concentrated in localized structural variants, repeat compartments, gene-family remodeling, and tandem arrays, is more consistent with ethnovariety-level differentiation within *B. caapi* than with species-level separation. This interpretation is further supported by the contrast between the strong collinearity observed between the caupuri and tucunacá assemblies and the more fragmented collinear relationships recovered in comparisons involving Salicaceae genomes, which represent deeper interspecific divergence within Malpighiales. However, formal taxonomic inference will require broader population sampling and dedicated species-delimitation analyses.

Because this study generated one chromosome-scale assembly per morphotype, the observed differences should be interpreted as genomic contrasts between the sampled caupuri and tucunacá accessions and as candidate morphotype-associated patterns, rather than as fixed genomic features of each morphotype. This sampling design does not reduce the value of the assemblies as high-quality genomic resources, but it defines the scope of inference. Broader population sampling, tissue-specific transcriptomics, anatomical analyses of stem development, and functional validation will be needed to determine which candidate genes, LARD-associated regions, and tandem arrays are consistently associated with stem-architecture variation in *B. caapi*.

## 5. Conclusions

Here, the chromosome-scale genomes of the caupuri and tucunacá ethnovarieties provide the first large-scale nuclear genomic resource for *B. caapi*, the principal liana component of ayahuasca. Ayahuasca has long been embedded in Indigenous and mestizo (caboclo) healing traditions across the Amazon, where it is used to address illness and mediate relationships between humans, plants, animals, and other-than-human beings [1, 3, 5, 96]. These longstanding traditions, together with the broader therapeutic, spiritual, and artistic contexts in which ayahuasca is now encountered [5, 97, 98] place *B. caapi* within a cultural history that extends beyond its pharmacological properties. This cultural significance adds broader relevance to the genomic resource presented here and situates the study within broader discussions on the dialogue between traditional knowledge and science [99]. Moreover, biomedical research has provided evidence for rapid antidepressant effects of ayahuasca in patients with recurrent depression [100, 101] and treatment-resistant depression [102], increasing interest in its therapeutic applications.

The two assemblies share a conserved chromosome-scale organization and paleopolyploid background within the limited genomic sampling currently available for Malpighiaceae, while their differences are concentrated in structural variation, repeat composition, gene-family copy number, assembly-restricted clusters, and comparatively recent tandem expansions. An indirect estimate placing the common node of the sampled accessions at approximately 1.13 Ma provides temporal context for recent repeat turnover and gene-family changes. The strongest candidate morphotype-biased signals involve TE-associated structural variants and tandem-expanded genes related to signaling, transport, hormone response, cell-wall organization, vascular and secondary-wall biology, and stem or shoot development.

The candidate loci summarized in Table 3 provide testable targets for studies of stem-architecture variation and morphotype-associated traits in *B. caapi*. By adding chromosome-scale genomic resources for a culturally significant tropical species, this study also contributes to reducing the genomic sampling gap for tropical plants [103] and extends comparative genomic resources for Malpighiaceae. The knowledge and practices of Indigenous and local communities associated with *B. caapi* and ayahuasca should be recognized as intellectual and cultural heritage, and research or commercial activities involving genetic resources or associated traditional knowledge should respect applicable principles of fair and equitable benefit-sharing [104].

## 6. List of abbreviations

AEK: ancestral eudicot karyotype
BLAST: Basic Local Alignment Search Tool
BUSCO: Benchmarking Universal Single-Copy Orthologs
CAFE5: Computational Analysis of gene Family Evolution, version 5
CDS: coding sequence
COG: Clusters of Orthologous Genes
CV: coefficient of variation
EC: Enzyme Commission
FDR: false discovery rate;
GO: Gene Ontology
Hi-C: high-throughput chromosome conformation capture
HiFi: high-fidelity
HPD: highest posterior density
Iso-Seq: isoform sequencing
ITS: internal transcribed spacer
Ka: nonsynonymous substitution rate
Ks: synonymous substitution rate
LARD: large retrotransposon derivative
LAI: LTR Assembly Index
LINE: long interspersed nuclear element
LTR: long terminal repeat
MCMCTree: Bayesian molecular-clock module of PAML
miRNA: microRNA
ncRNA: non-coding RNA
NOR: nucleolar organizer region
OG: orthogroup
OMArk: orthology-based genome annotation quality assessment
PAML: Phylogenetic Analysis by Maximum Likelihood
PASA: Program to Assemble Spliced Alignments
QV: quality value
RNA-seq: RNA sequencing
rRNA: ribosomal RNA
snoRNA: small nucleolar RNA
snRNA: small nuclear RNA
SRA: Sequence Read Archive
SRP RNA: signal recognition particle RNA
SV: structural variant
TPM: Transcripts Per Million
TE: transposable element
TIR: terminal inverted repeat
TPM: transcripts per million
tRNA: transfer RNA
WGD: whole-genome duplication
WGDI: Whole-Genome Duplication Integrated analysis

## 7. Declarations

### 7.1. Availability of data and materials

Access to the genetic resources and associated traditional knowledge was conducted in accordance with applicable Brazilian regulations on access and benefit-sharing and was registered in SisGen under registration number #A2A72C6. The raw sequencing data have been deposited in the NCBI Sequence Read Archive under BioProject accessions PRJNA1497017 for the caupuri ethnovariety and PRJNA997335 for the tucunacá ethnovariety. The associated SRA accessions are SRR39699010 for caupuri HiFi genomic DNA sequencing, SRR39699011 for caupuri Iso-Seq data, SRR39697887 for tucunacá HiFi genomic DNA sequencing, and SRR39697886 for caupuri Hi-C data. Genome assembly and annotation files are also available through the Plant Genomics database (https://plantgenomics.ncc.unesp.br/Genomes/index.php?id=Tucunaca and https://plantgenomics.ncc.unesp.br/Genomes/index.php?id=Caupuri) and the Zenodo repository under DOI/accession https://doi.org/10.5281/zenodo.21725632. The caupuri whole-genome shotgun project has been deposited at DDBJ/ENA/GenBank under the accession PRJNA1497017 and Tucunacá is under the accession PRJNA997335.

### 7.2. Additional files

Additional file 1 (.pdf). Supplementary figures and small supplementary tables. Contains Figures S1–S9 and Tables S1–S5, S8, and S19, together with the legends of Tables S6–S7 and S9–S18. Additional file 2 (.xlsx). Large supplementary tables. Contains Tables S6–S7 and S9– S18.

### 7.3. Competing interests

The authors declare no conflict of interest.

### 7.4. Funding

This study was financed by Fundação de Amparo à Pesquisa do Estado do Rio de Janeiro – FAPERJ grants CNE E-26/ 202.780/2018 and E-26/200.940/2022 to FP; and also, by Fundação de Amparo à Pesquisa do Estado do São Paulo – FAPESP grants #2023/10314-4 to VT, #2022/15735-5 to AMV, #2022/05890-3 to APM, and #2021/10639-5 to VFOM. VT, SRS, VFOM, APM, and AMV are affiliated with the Center for Research on Biodiversity Dynamics and Climate Change (CEPID-FAPESP, grant #2021/10639-5). FP, AMV, VFOM, and APM are recipients of Researcher Fellowships from Conselho Nacional de Desenvolvimento Científico e Tecnológico (CNPq). Finally, this work was supported by Fundação Araucária (NAPI Bioinformática, #66.2021) and CNPq (#440412/2022-6) to ARP. These funding agencies had no role in study design, data collection, analysis, interpretation, or manuscript writing.

### 7.5. Authors’ contributions

Conceptualization: VT, FP, AMV. Data curation: VT, FP, AMV.

Formal analysis: VT, FP, PCB, APM, MCA, ARP, SRS, VFOM, AMV.

Funding acquisition: FP, VFOM, AMV.

Investigation: FP, APM, SRS, VFOM, JBFB, DRO, SSSL, AMV.

Project administration: FP, AMV.

Resources: FP, APM, SRS, VFOM, JBFB, SSSL, AMV.

Supervision: FP, AMV.

Validation: VT, FP, PCB, MCA, ARP, AMV.

Visualization: VT, APM, MCA, ARP, AMV. Writing – original draft: VT, FP, AMV.

Writing – review & editing: VT, FP, APM, ARP, SRS, VFOM, JBFB, DRO, SSSL, AMV.

All authors read and approved the final manuscript.

## 7.6. Acknowledgments

We thank the students and researchers Sérgio Schuaba, Rafael Andrade, João Gisbert, Eduarda Tavares, Henrique Pinheiro, and Maria Alice Corrêa, who contributed to the initial analysis in this study. We are also grateful to the “Centro Espírita Beneficente União do Vegetal” (CEBUDV) for providing the two ethnovarieties and for supporting this research. We are deeply grateful to Mestre Ricardo Teixeira and the members of Núcleo Menino Galante for their assistance and traditional knowledge. We acknowledge the long-standing knowledge, practices, and stewardship of Indigenous and local communities associated with *Banisteriopsis caapi* and the ayahuasca brew and recognize these traditions as an important part of their cultural and intellectual heritage. Research involving these biological resources and associated knowledge should be conducted with respect to the communities from which they originate and for applicable principles of access and fair and equitable benefit-sharing. We also thank the Arizona Genomics Institute (Tucson, AZ, USA) for performing all the HMW DNA and mRNA extraction and sequencing.

## Notes

https://zenodo.org/records/21725632

### Competing Interest Statement

The authors have declared no competing interest.

