## Additional File 1 for "Comparative chromosome-scale genomics of the ayahuasca liana *Banisteriopsis caapi* reveals paleopolyploid ancestry and recent tandem-gene expansions as candidate signatures of ethnovariety differentiation"

**Supplementary Information**

**Supplementary Figures**

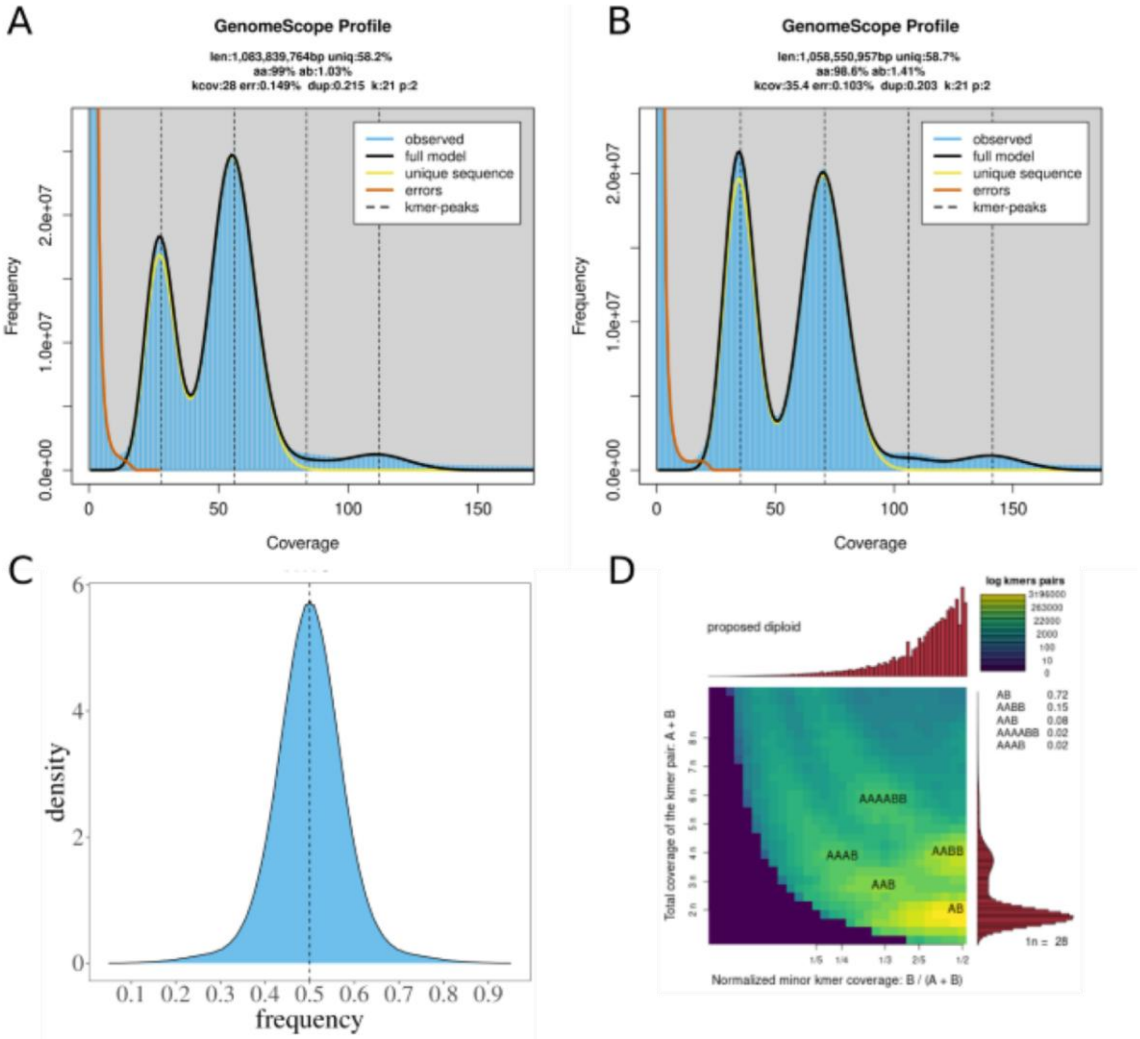


**Figure S1. K-mer-based genome profiling and ploidy assessment of the *Banisteriopsis caapi* morphotypes**

(A–B) GenomeScope profiles generated from k-mer spectra for tucunacá (A) and caupuri (B). (C) Representative PloidyFrost allele-frequency profile used for ploidy inference. (D) Representative Smudgeplot k-mer pair profile used for ploidy inference. PloidyFrost and Smudgeplot profiles are shown as representative outputs because the same ploidy pattern was inferred for both morphotypes.

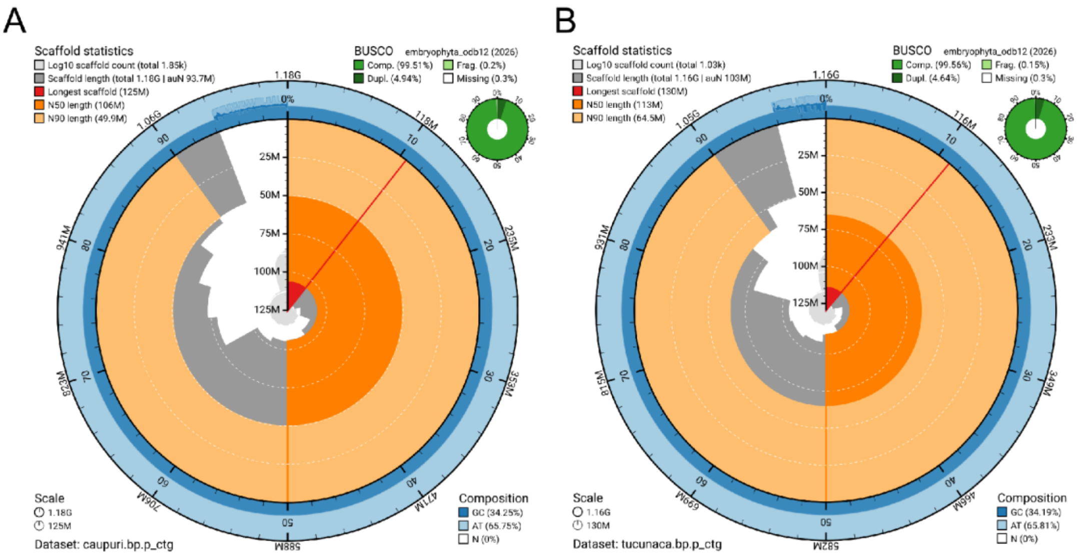


**Figure S2. Snail-plot summaries of the primary draft genome assemblies of the caupuri and tucunacá ethnovarieties of *Banisteriopsis caapi***

Snail plots summarize assembly contiguity, scaffold-length distribution, nucleotide composition, and BUSCO completeness for the primary draft assemblies generated before chromosome-scale scaffolding. (A) Caupuri draft assembly. (B) Tucunacá draft assembly. The circumference represents the cumulative assembly span, while scaffolds are arranged in decreasing order of length. The central gray areas represent the size-ordered scaffold-length distribution and the log-scaled cumulative scaffold count. Red, dark-orange, and light-orange segments indicate the longest scaffold, scaffold N50, and scaffold N90, respectively. The outer blue track shows GC and AT composition across the assembly, and the inset charts report complete, duplicated, fragmented, and missing BUSCOs based on the embryophyta_odb12 dataset. Total assembly span, scaffold count, auN, longest-scaffold length, N50, and N90 are reported within each panel. Snail plots were generated with BlobTK.

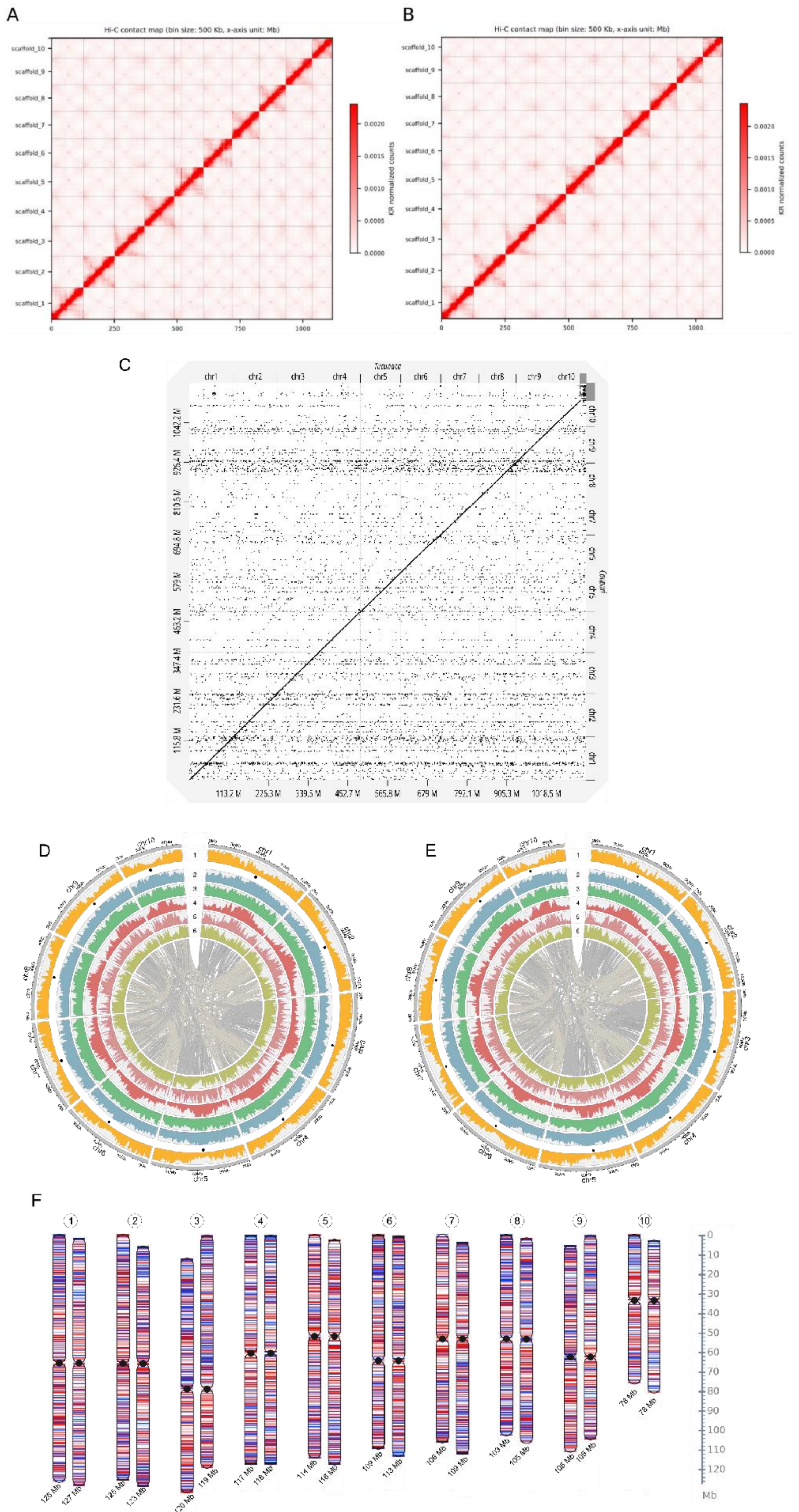
A
B
Hi-C contact map (bin size: 500 Kb, x-axis unit: Mb)
Hi-C contact map (bin size: 500 Kb, x-axis unit: Mb)
scaffold_10
scaffold_9
scaffold_8
scaffold_7
scaffold_6
scaffold_5
scaffold_4
scaffold_3
scaffold_2
scaffold_1
KR normalized counts
0.0020
0.0015
0.0010
0.0005
0.0000
0
250
500
750
1000
C
chr1
chr2
chr3
chr4
chr5
chr6
chr7
chr8
chr9
chr10
113.2 M
226.3 M
339.5 M
452.7 M
565.8 M
679 M
792.1 M
905.3 M
1018.5 M
D
E
F
Mb

**Figure S3. Genome assembly and annotation overview of the caupuri and tucunacá assemblies**

(A–B) Hi-C contact maps of the caupuri (A) and tucunacá (B) chromosome-scale assemblies. Hi-C data were generated only for the caupuri morphotype. To avoid possible assembly errors during the scaffolding of the tucunacá morphotype, the Hi-C reads were not employed to perform self-correction on the contigs assembled before scaffolding. For the caupuri morphotype, the scaffolding was performed with default parameters. (C) Dot-plot comparison via the dgenies tool (https://dgenies.toulouse.inra.fr/) between the tucunacá (X axis) and caupuri (Y axis) morphotypes. The results indicate that only ~12% of both genomes lack matches. (D–E) Circular genome maps of the caupuri and tucunacá chromosome-scale assemblies. From the inner to the outer circle: (1) genes, (2) LTR Copia, (3) LTR Gypsy, (4) LINE, (5) TIR, and (6) Helitron. Black dots represent the centromere positions. (F) Chromosome-scale representation of the two morphotype assemblies, showing the ten corresponding chromosomes, chromosome lengths, and annotated chromosomal features.

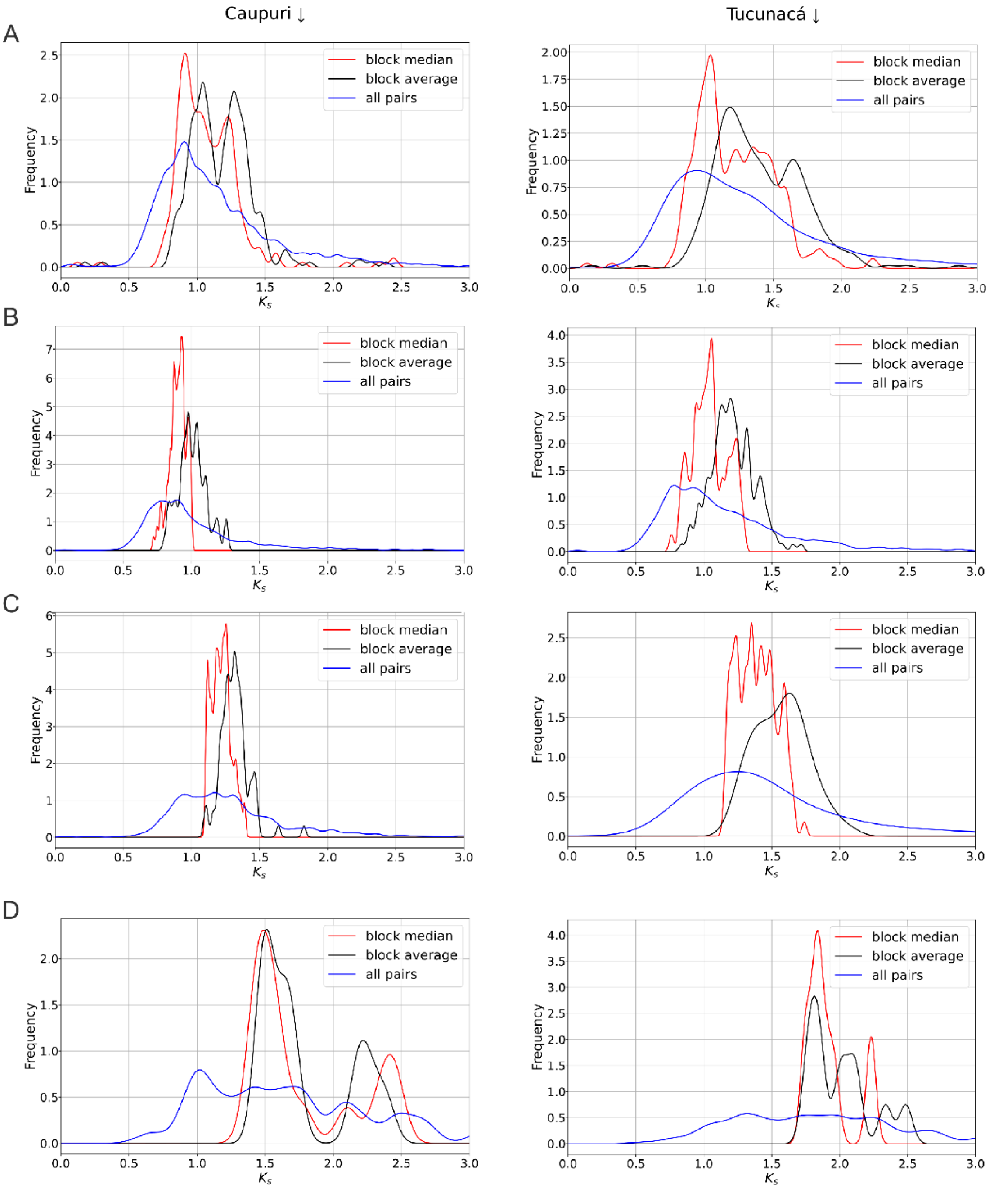


**Figure S4. Ks distributions of duplicated gene pairs in the caupuri and tucunacá morphotypes**

Ks density plots for duplicated gene pairs in the caupuri and tucunacá genomes. Panels show distributions calculated using block median, block average, and all-pair estimates with WGDI. Panel (A) shows the complete Ks profile, whereas panels (B-D) show selected ranges of 0.50-1.00 (B), 1.10-1.40 (C), and 1.4-3.0 (D). The plots display Ks values on the x-axis and frequencies on the y-axis.

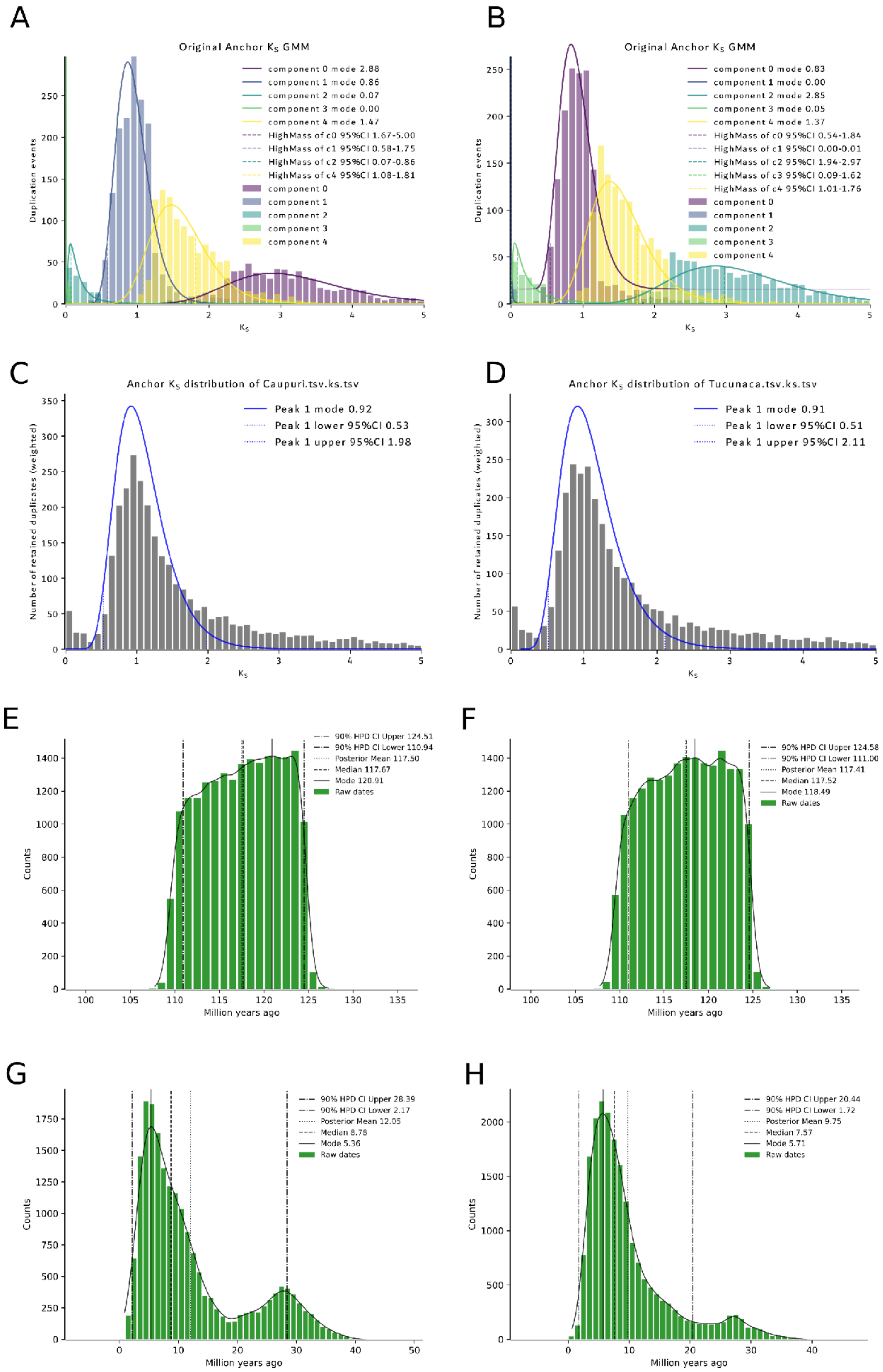
A
Original Anchor $K_S$ GMM
component 0 mode 2.88
component 1 mode 0.86
component 2 mode 0.07
component 3 mode 0.00
component 4 mode 1.47
HighMass of c0 95%CI 1.67-5.00
HighMass of c1 95%CI 0.58-1.75
HighMass of c2 95%CI 0.07-0.86
HighMass of c4 95%CI 1.08-1.81
Duplication events
B
Original Anchor $K_S$ GMM
component 0 mode 0.83
component 1 mode 0.00
component 2 mode 2.85
component 3 mode 0.05
component 4 mode 1.37
HighMass of c0 95%CI 0.54-1.84
HighMass of c1 95%CI 0.00-0.01
HighMass of c2 95%CI 1.94-2.97
HighMass of c3 95%CI 0.09-1.62
HighMass of c4 95%CI 1.01-1.76
C
Anchor $K_S$ distribution of Caupuri.tsv.ks.tsv
Peak 1 mode 0.92
Peak 1 lower 95%CI 0.53
Peak 1 upper 95%CI 1.98
Number of retained duplicates (weighted)
D
Anchor $K_S$ distribution of Tucunaca.tsv.ks.tsv
Peak 1 mode 0.91
Peak 1 lower 95%CI 0.51
Peak 1 upper 95%CI 2.11
E
90% HPD CI Upper 124.51
90% HPD CI Lower 110.94
Posterior Mean 117.50
Median 117.67
Mode 120.91
Raw dates
Counts
Million years ago
F
90% HPD CI Upper 124.58
90% HPD CI Lower 111.00
Posterior Mean 117.41
Median 117.52
Mode 118.49
Raw dates
G
90% HPD CI Upper 28.39
90% HPD CI Lower 2.17
Posterior Mean 12.05
Median 8.78
Mode 5.36
Raw dates
H
90% HPD CI Upper 20.44
90% HPD CI Lower 1.72
Posterior Mean 9.75
Median 7.57
Mode 5.71
Raw dates

**Figure S5. Whole-genome duplication analysis and duplication-time estimates for the caupuri and tucunacá morphotypes**

WGD_v2 analyses of duplicated gene pairs in the caupuri (A and C) and tucunacá genomes (B and D). Panels include Gaussian mixture models fitted to anchor-pair Ks distributions (A and B), anchor Ks distributions with inferred peaks (C and D). Stars represent WGD events. (E-H) duplication-time distributions estimated from Ks-derived age intervals peaks.

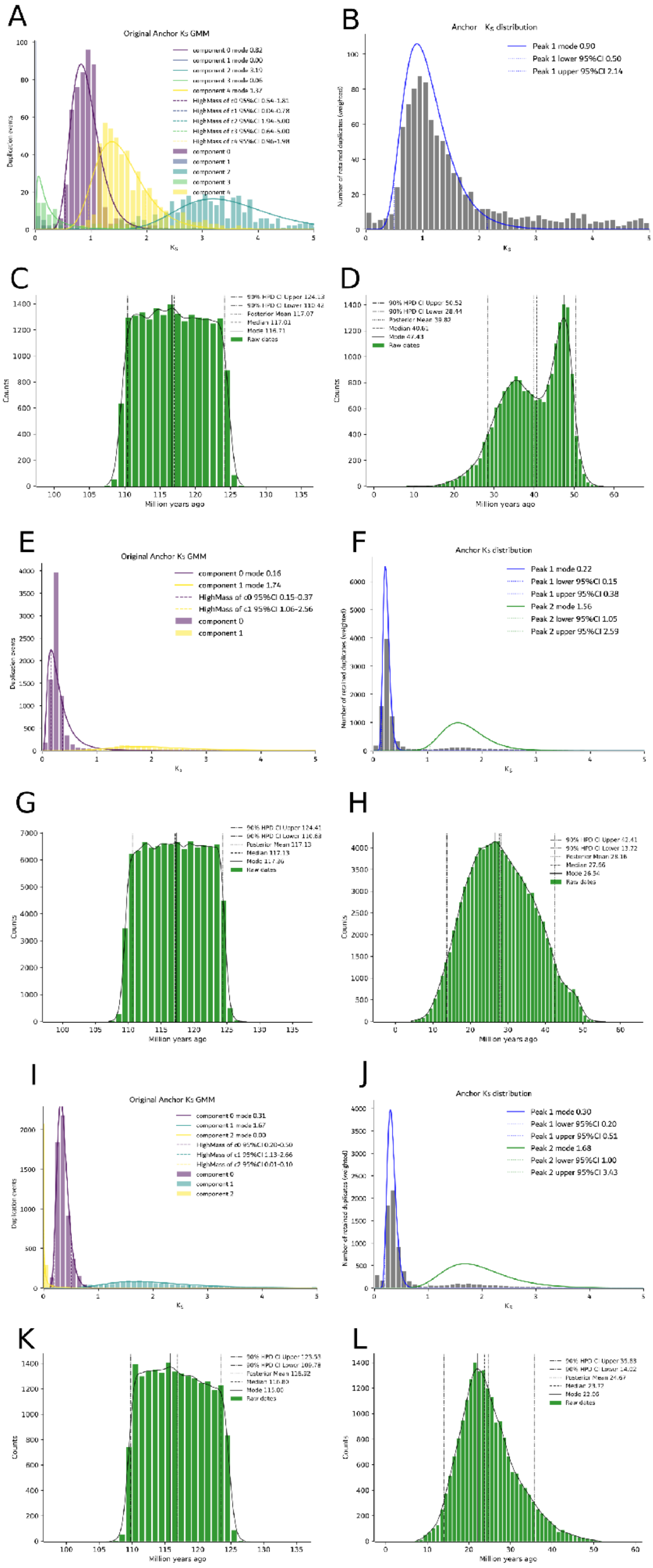
A
Original Anchor Ks GMM
B
Anchor Ks distribution
Peak 1 mode 0.90
Peak 1 lower 95%CI 0.50
Peak 1 upper 95%CI 2.14
C
Million years ago
Counts
D
Million years ago
Counts
E
Original Anchor Ks GMM
component 0 mode 0.16
component 1 mode 1.74
HighMass of c0 95%CI 0.15-0.37
HighMass of c1 95%CI 1.06-2.56
component 0
component 1
Duplication events
F
Anchor Ks distribution
Peak 1 mode 0.22
Peak 1 lower 95%CI 0.15
Peak 1 upper 95%CI 0.38
Peak 2 mode 1.56
Peak 2 lower 95%CI 1.05
Peak 2 upper 95%CI 2.59
G
Million years ago
Counts
H
Million years ago
Counts
I
Original Anchor Ks GMM
J
Anchor Ks distribution
Peak 1 mode 0.30
Peak 1 lower 95%CI 0.20
Peak 1 upper 95%CI 0.51
Peak 2 mode 1.68
Peak 2 lower 95%CI 1.00
Peak 2 upper 95%CI 3.43
K
Million years ago
Counts
L
Million years ago
Counts

**Figure S6. Comparative whole-genome duplication analyses in *Malpighia emarginata* (A-D), *Populus trichocarpa* (E-H), and *Salix purpurea* (I-L)**

WGD_v2 analyses of duplicated gene pairs in the comparative genomes used to contextualize the *B. caapi* results. Panels show Gaussian mixture models fitted to anchor-pair Ks distributions, anchor Ks distributions with inferred peaks, and duplication-time distributions for each sampled genome.

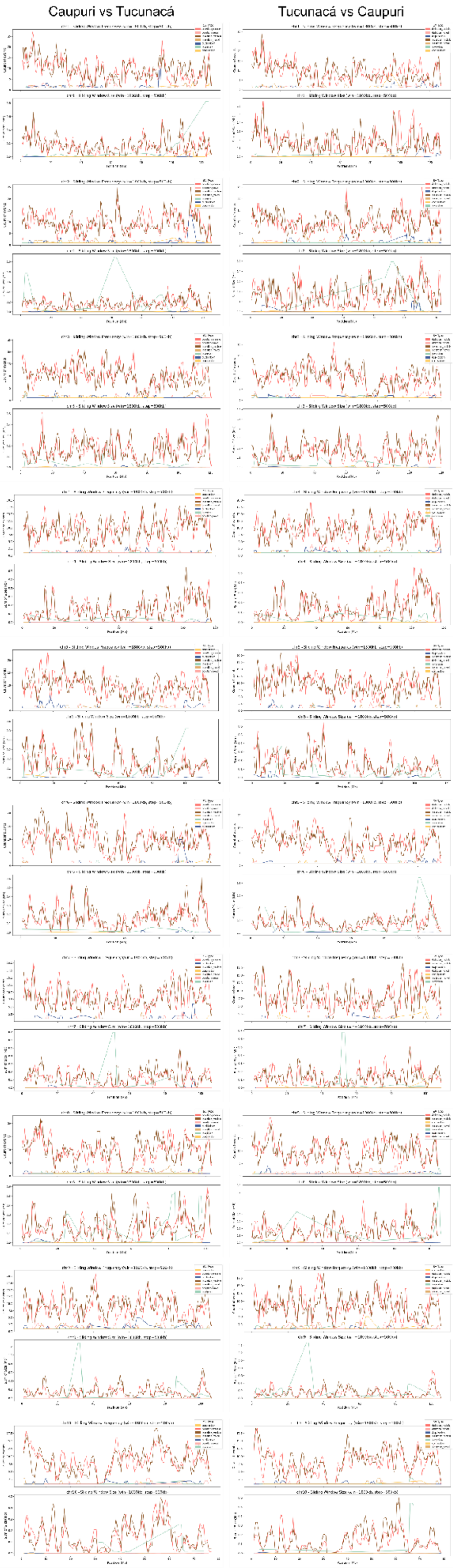
Caupuri vs Tucunacá
Tucunacá vs Caupuri

**Figure S7. Chromosome-wise structural variation profiles between caupuri and tucunacá**
Structural variant distributions detected in reciprocal whole-genome comparisons between the caupuri and tucunacá assemblies. Panels show chromosome-wise SV span and SV frequency for mobile element-associated deletions and insertions, novel deletions and insertions, duplications, inversions, and contractions.

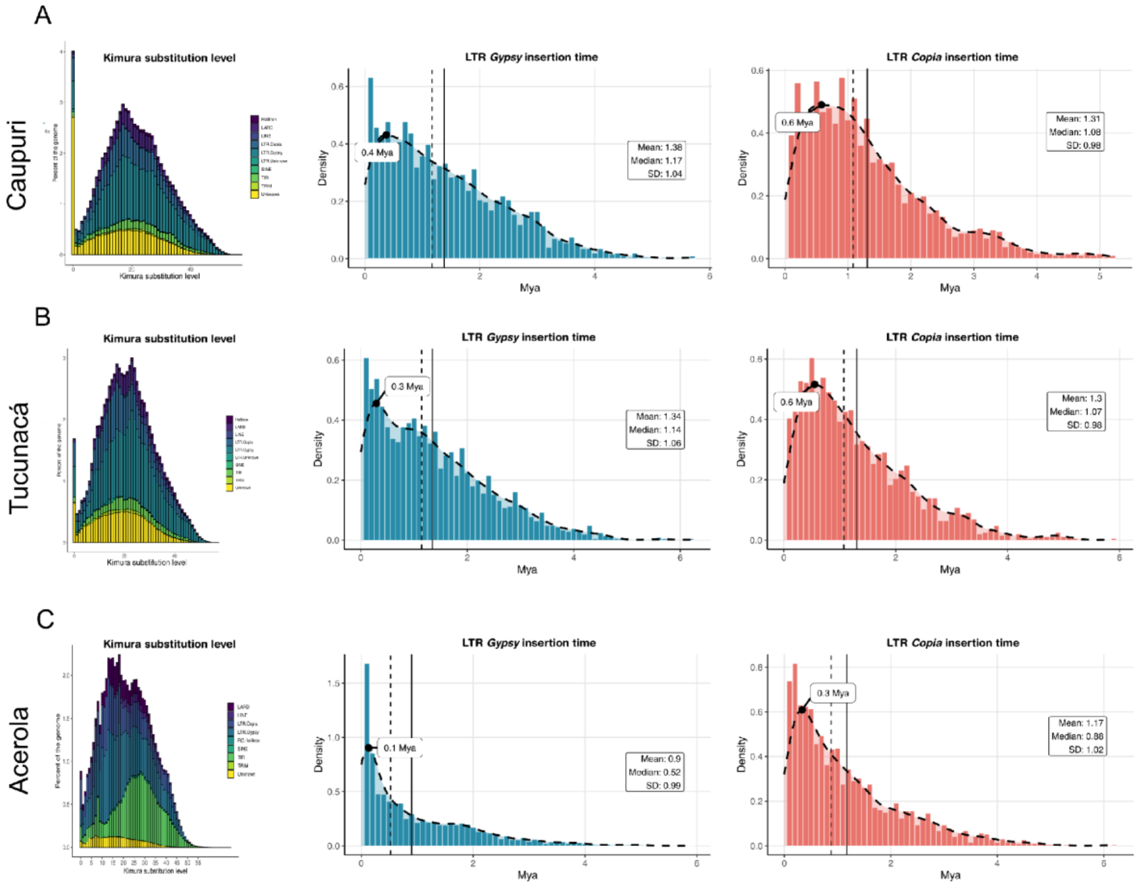


**Figure S8. Repeat divergence landscapes and intact LTR insertion-time distributions in caupuri, tucunacá, and acerola (*M. emarginata)***

Kimura divergence landscapes and intact LTR insertion-time distributions for caupuri, tucunacá, and acerola. For each genome, the left panel shows the Kimura divergence distribution of major repeat classes, the middle panel shows LTR/*Gypsy* insertion-time estimates, and the right panel shows LTR/*Copia* insertion-time estimates. Insertion times are shown in millions of years ago.

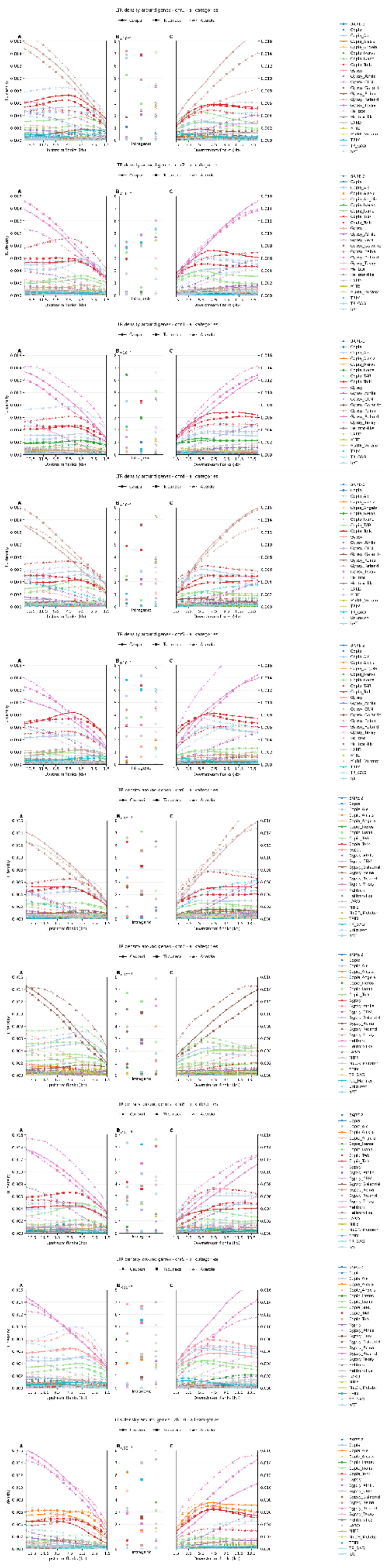

**Figure S9. Chromosome-resolved TE-density profiles around genes in caupuri, tucunacá, and acerola**

Per-chromosome TE-density profiles around annotated genes for all selected TE categories in caupuri, tucunacá, and acerola. For each chromosome, upstream flanking regions, intragenic regions, and downstream flanking regions are shown separately. Lines represent TE-density profiles for individual TE categories, with the three genomes displayed together for comparison.

### Supplementary Tables

**Table S1. Sporophytic genome size (2C) estimation by flow cytometry from two different morphotypes of *Banisteriopsis caapi*, caupuri and tucunacá**

Raw flow-cytometry measurements are shown for each analyzed individual in technical triplicate, together with the coefficient of variation (CV%) of each sample and of the internal reference standard, *Raphanus sativus* L. 'Saxa' (2C = 1.11 pg). Morphotype-level averages and standard deviations are reported for descriptive purposes. For statistical comparison between morphotypes, technical triplicates were averaged within each individual, and individual mean 2C values were compared using Welch's two-sample t-test.

| Morphotype | Individual | Repetition | 2C (pg) | Sample CV (%) | Internal reference standard CV (%) | SD |
|---|---|---|---|---|---|---|
| Caupuri | 1† | A | 2.740 | 4.05 | 3.79 | |
| | | B | 2.744 | 4.425 | 3.61 | |
| | | C | 2.70 | 5.435 | 3.93 | |
| | 2 | A | 2.62 | 3.12 | 2.06 | |
| | | B | 2.60 | 3.60 | 2.73 | |
| | | C | 2.58 | 3.54 | 2.46 | |
| | **Average** | | **2.66** | **4.02** | **3.10** | **0.07 (2.70%)** |
| Tucunacá | 1† | A | 2.69 | 4.84 | 3.10 | |
| | | B | 2.66 | 4.03 | 2.76 | |
| | | C | 2.70 | 4.24 | 2.74 | |
| | 2 | A | 2.70 | 3.92 | 2.96 | |
| | | B | 2.79 | 4.27 | 3.02 | |
| | | C | 2.71 | 3.89 | 3.21 | |
| | 3 | A | 2.71 | 3.57 | 2.65 | |
| | | B | 2.75 | 3.13 | 3.37 | |
| | | C | 2.67 | 3.18 | 3.63 | |
| | **Average** | | **2.70** | **3.90** | **3.09** | **0.05 (1.72%)** |
| | **Average** | | **2.69** | **3.95** | **3.09** | **0.05 (2.03%)** |

Welch's two-sample t-test based on individual mean 2C values: $t = -0.684$, $df = 1.10$, $p = 0.609$.
† Individuals submitted to DNA sequencing.

**Table S2. Final genome assembly statistics and completeness scores of the *Banisteriopsis caapi* (mariri), two different morphotypes, caupuri and tucunacá**

Summary of assembly contiguity, scaffold statistics, BUSCO genome and annotation completeness, OMArk annotation assessment, and LTR Assembly Index values for the two *B. caapi* morphotype assemblies.

| | Caupuri | Tucunacá |
|---|---|---|
| Genome assembly size | 1,158,016,917 | 1,131,625,749 |
| Number of scaffolds | 1,528 | 292 |
| Longest scaffold | 126,276,660 | 127,231,723 |
| Shortest scaffold | 14,733 | 11,444 |
| Number of scaffolds > 1K | 1,528 | 292 |
| Number of scaffolds > 10K | 1,528 | 292 |
| Number of scaffolds > 100K | 32 | 28 |
| Number of scaffolds > 1M | 10 | 11 |
| Number of scaffolds > 10M | 10 | 10 |
| Mean scaffold size | 757,864 | 3,875,431 |
| Median scaffold size | 31,764 | 41,170 |
| N50 scaffold length | 114,016,100 | 115,587,645 |
| L50 scaffold count | 5 | 5 |
| BUSCO† genome | C:99.3%[S:94.3%,D:5.0%], F:0.2%,M:0.4%,n:2026 | C:99.6%[S:94.9%,D:4.7%], F:0.1%,M:0.2%,n:2026 |
| BUSCO† annotation | C:98.8%[S:94.4%,D:4.3%], F:0.7%,M:0.5%,n:2026 | C:99.0%[S:94.7%,D:4.3%], F:0.5%,M:0.5%,n:2026 |
| OMArk‡ | A:86.01%[P:18.19%,F:8.50%], I:2.52%[P:1.36%,F:0.27%], C:0.00%[P:0.00%,F:0.00%], U:11.48% | A:85.09%[P:19.66%,F:8.76%], I:2.44%[P:1.52%,F:0.28%], C:0.00%[P:0.00%,F:0.00%], U:12.46% |
| LAI | 13.88 | 14.05 |

†BUSCO classification: C: complete; S: complete and single copy; D: complete and duplicated; F: fragmented; M: missing; n: number of BUSCO groups searched; embryophyta_odb12 dataset were employed.

‡OMArk classification: A: taxonomically consistent; P: partial hits; F: fragmented; I: taxonomically inconsistent; C: likely contamination; U: unknown. Rosids clade (NCBI:txid71275) was employed in OMArk processing, which contains 10,030 HOGs (hierarchical orthologous groups).

**Table S3. Predicted centromere coordinates in the caupuri and tucunacá chromosome-scale assemblies**

Chromosome-wise coordinates of predicted centromeric regions in the ten chromosome-scale scaffolds of caupuri and tucunacá. Centromere start, end, and length for each chromosome are distributed in columns and are reported in base pairs.

| **Morphotypes** | **Caupuri** | | | **Tucunacá** | | |
|---|---|---|---|---|---|---|
| Chromosomes | Start | End | Length | Start | End | Length |
| chr1 | 65,516,559 | 65,546,559 | 30,000 | 64,169,390 | 64,199,390 | 30,000 |
| chr2 | 65,801,989 | 65,831,989 | 30,000 | 60,214,254 | 60,244,254 | 30,000 |
| chr3 | 66,638,289 | 66,668,289 | 30,000 | 78,924,695 | 78,954,695 | 30,000 |
| chr4 | 60,219,734 | 60,249,734 | 30,000 | 60,455,160 | 60,485,160 | 30,000 |
| chr5 | 51,983,776 | 52,013,776 | 30,000 | 49,677,939 | 49,707,939 | 30,000 |
| chr6 | 64,365,054 | 64,395,054 | 30,000 | 64,092,045 | 64,122,045 | 30,000 |
| chr7 | 53,232,094 | 53,262,094 | 30,000 | 49,657,759 | 49,687,759 | 30,000 |
| chr8 | 53,198,018 | 53,228,018 | 30,000 | 51,884,744 | 51,914,744 | 30,000 |
| chr9 | 56,967,848 | 56,997,848 | 30,000 | 62,456,711 | 62,486,711 | 30,000 |
| chr10 | 33,656,195 | 33,686,195 | 30,000 | 30,986,801 | 31,016,801 | 30,000 |

**Table S4. Predicted telomeric repeat distribution in the caupuri and tucunacá chromosome-scale assemblies**

Chromosome-wise telomere status and telomeric repeat counts at the left and right ends of the ten chromosome-scale scaffolds of caupuri and tucunacá.

| | **Caupuri** | | | **Tucunacá** | | |
|---|---|---|---|---|---|---|
| Chr ID | Status | Left num | Right num | Status | Left num | Right num |
| chr1 | left | 929 | 0 | left | 2631 | 0 |
| chr2 | both | 482 | 1,195 | both | 1826 | 1,560 |
| chr3 | left | 1404 | 0 | left | 2122 | 0 |
| chr4 | both | 1161 | 382 | both | 579 | 2,863 |
| chr5 | both | 447 | 781 | both | 1673 | 1,658 |
| chr6 | both | 436 | 457 | both | 2080 | 2,347 |
| chr7 | right | 0 | 957 | both | 2105 | 2,253 |
| chr8 | left | 643 | 0 | left | 2446 | 0 |
| chr9 | left | 342 | 0 | left | 2619 | 0 |
| chr10 | left | 719 | 0 | left | 762 | 0 |

**Table S5. Genome annotation summary features and statistics for the *Banisteriopsis caapi* (mariri) two different morphotypes, caupuri and tucunacá, complete genome assemblies**

| Features | Caupuri | Tucunacá |
|---|---|---|
| - Number of genes | 59,836 | 47,458 |
| -- Number of exons | 404,027 | 405,358 |
| -- Number of introns | 322,283 | 333,802 |
| -- Number of CDSs | 81,744 | 71,556 |
| --- complete CDS | 81,207 | 71,318 |
| --- start, no stop CDS | 289 | 116 |
| --- stop, no start CDS | 214 | 91 |
| --- no stop, no start CDS | 34 | 31 |
| -- Overlapping genes | 6,725 | 5,402 |
| -- Contained genes | 1,955 | 1,262 |
| -- Shortest gene | 153 | 150 |
| -- Longest gene | 208,917 | 276,605 |
| - Longest CDS | 15,246 | 16,284 |
| - mean gene length | 2,566 | 2,814 |
| - mean CDS length | 1,194 | 1,265 |
| - mean exons per gene | 5 | 6 |
| - mean introns per gene | 4 | 5 |
| - WGD gene pairs | 16,345 | 11,717 |
| - Tandem gene pairs | 5,074 | 5,062 |
| - Proximal duplicated gene pairs | 6,694 | 6,634 |
| - Dispersed duplicated gene pairs | 11,686 | 9,264 |
| - Singletons | 3,431 | 3,403 |
| - Transposed gene pairs | 9,085 | 6,126 |
| - Retrocopies | 1,669 | 1,110 |
| -- Parental Gene | 497 | 441 |
| --- Chimerical | 1,004 | 623 |
| --- Pseudogene | 246 | 189 |
| --- Retrogene | 419 | 298 |
| - % of genome covered by genes | 13.3 | 11.8 |
| - % of genome covered by CDS | 8.4 | 8.0 |
| - % of genome covered by TEs | 77.96% | 78.27% |
| -- % of Class I Elements | 55.47% | 57.24% |
| --- % of LTR Gypsy | 28.39% | 28.33% |
| --- % of LTR Copia | 13.30% | 15.07% |
| --- % of LTR non-autonomous | 10.43% | 9.52% |
| -- % of Class II Elements | 7.69% | 8.28% |
| --- % TIRs | 6.08% | 6.47% |
| --- % Helitron | 1.61% | 1.80% |

**Table S6. Complete structural and Functional annotation of *Banisteriopsis caapi***
Gene-level annotation of the predicted gene sets for caupuri (A) and tucunacá (B), including functional descriptions, genomic coordinates, duplication topology, COG, EC, GO terms, and TPM.

***Provided as an external Excel file***

**Table S7. Genome-wide non-coding RNA annotation summary for the caupuri and tucunacá ethnovarieties of *Banisteriopsis caapi* and *Malpighia emarginata***
Numbers of chromosome-anchored non-coding RNA loci assigned to tRNA, snoRNA, rRNA, snRNA, miRNA, signal recognition particle RNA (SRP RNA), ribozyme, RNase P/MRP RNA, lncRNA, and other RNA classes. Totals represent non-redundant loci retained on chromosome-scale sequences after filtering and merging.

***Provided as an external Excel file***

**Table S8. Chromosome-wise distribution, frequency, and cumulative size of structural variations (SVs) in *Banisteriopsis caapi* caupuri and tucunacá morphotype**

Summary of structural variants detected in reciprocal whole-genome comparisons between the caupuri and tucunacá assemblies. The table reports, by chromosome and SV class, the number of events and cumulative genomic span for contractions, mobile element-associated deletions and insertions, novel deletions and insertions, duplications, inversions, and translocations. Subtables show SV frequencies and total SV span separately for each morphotype.

**Table S8A. Caupuri: Frequency of Structural Variation Types by Chromosome**

| Chromosome | Contraction | Deletion_mobile | Deletion_novel | Duplication | Insertion_mobile | Insertion_novel | Inversion | Translocation | TOTAL |
|---|---|---|---|---|---|---|---|---|---|
| 1 | 14 | 637 | 43 | 42 | 713 | 24 | 14 | 50 | 1,537 |
| 2 | 19 | 684 | 38 | 46 | 700 | 31 | 9 | 41 | 1,568 |
| 3 | 8 | 709 | 37 | 39 | 690 | 49 | 17 | 56 | 1,605 |
| 4 | 17 | 602 | 27 | 38 | 642 | 39 | 9 | 49 | 1,423 |
| 5 | 3 | 618 | 32 | 51 | 645 | 28 | 12 | 33 | 1,422 |
| 6 | 16 | 582 | 29 | 33 | 658 | 20 | 11 | 45 | 1,394 |
| 7 | 9 | 498 | 29 | 37 | 494 | 22 | 8 | 65 | 1,162 |
| 8 | 5 | 548 | 32 | 27 | 606 | 30 | 18 | 37 | 1,303 |
| 9 | 6 | 494 | 49 | 39 | 569 | 34 | 11 | 52 | 1,254 |
| 10 | 4 | 378 | 19 | 19 | 372 | 22 | 9 | 23 | 846 |
| **TOTAL** | 101 | 5,750 | 335 | 371 | 6,089 | 299 | 118 | 451 | 13,514 |

**Table S8B. Caupuri: Total Size of Structural Variations by Chromosome (bp)**

| Chromosome | Size (bp) | Contrac-tion | Deletion_ Mobile | Deletion_ Novel | Duplication | Insertion_ Mobile | Insertion_ Novel | Inversion | Translocation | TOTAL | % occupied |
|---|---|---|---|---|---|---|---|---|---|---|---|
| 1 | 126,236,216 | 154,335 | 11,075,049 | 13,510 | 116,721 | 13,574,163 | 17,701 | 1,342,805 | 230,493,929 | 26,294,284 | 20.83 |
| 2 | 125,311,022 | 268,692 | 11,349,059 | 12,561 | 70,073 | 12,420,731 | 7,172 | 2,692,250 | 241,848,070 | 26,820,538 | 21.40 |
| 3 | 119,580,670 | 10,844 | 11,183,553 | 9,284 | 75,710 | 10,848,660 | 9,461 | 651,922 | 229,257,542 | 22,789,434 | 19.06 |
| 4 | 116,948,394 | 4,109 | 9,275,001 | 8,349 | 82,105 | 11,249,343 | 16,069 | 263,035 | 219,895,641 | 20,898,011 | 17.87 |
| 5 | 114,004,802 | 33,641 | 10,179,718 | 8,680 | 82,239 | 11,315,116 | 8,867 | 1,318,600 | 196,993,576 | 22,946,861 | 20.13 |

| | | | | | | | | | | | |
|---|---|---|---|---|---|---|---|---|---|---|---|
| 6 | 109,297,113 | 36,575 | 10,358,200 | 9,170 | 112,707 | 11,436,058 | 5,389 | 263,102 | 211,143,007 | 22,221,201 | 20.33 |
| 7 | 106,097,994 | 2,473 | 7,391,300 | 9,989 | 201,531 | 8,864,468 | 6,254 | 934,984 | 203,963,898 | 17,410,999 | 16.41 |
| 8 | 102,529,633 | 962 | 8,979,223 | 7,916 | 72,714 | 10,898,094 | 7,488 | 2,123,275 | 173,777,910 | 22,089,672 | 21.54 |
| 9 | 105,477,578 | 1,731 | 7,697,014 | 13,335 | 50,606 | 10,042,916 | 9,893 | 2,015,772 | 193,170,437 | 19,831,267 | 18.80 |
| 10 | 76,276,120 | 931 | 6,193,309 | 4,793 | 17,367 | 6,391,729 | 4,775 | 443,278 | 145,770,755 | 13,056,182 | 17.12 |
| **TOTAL** | 1,101,759,542 | 514,293 | 93,681,426 | 97,587 | 881,773 | 107,041,278 | 93,069 | 12,049,023 | 2,046,314,765 | 214,358,449 | 19.46 |
| **% occupied** | | 0.05 | 8.50 | 0.01 | 0.08 | 9.72 | 0.01 | 1.09 | 185.73 | 19.46 | |

**Table S8C. Tucunacá: Frequency of Structural Variation Types by Chromosome**

| Chromosome | Contraction | Deletion_Mobile | Deletion_Novel | Duplication | Insertion_Mobile | Insertion_Novel | Inversion | Translocatio n | TOTAL |
|---|---|---|---|---|---|---|---|---|---|
| 1 | 17 | 675 | 24 | 46 | 667 | 44 | 8 | 35 | 1516 |
| 2 | 7 | 676 | 29 | 43 | 706 | 39 | 6 | 48 | 1554 |
| 3 | 10 | 656 | 51 | 26 | 750 | 35 | 14 | 63 | 1605 |
| 4 | 15 | 605 | 39 | 38 | 639 | 27 | 14 | 51 | 1428 |
| 5 | 10 | 627 | 29 | 33 | 638 | 29 | 22 | 27 | 1415 |
| 6 | 5 | 613 | 20 | 39 | 617 | 28 | 14 | 66 | 1402 |
| 7 | 20 | 475 | 23 | 23 | 530 | 29 | 12 | 30 | 1142 |
| 8 | 9 | 575 | 31 | 23 | 584 | 30 | 12 | 62 | 1326 |
| 9 | 18 | 509 | 35 | 25 | 524 | 50 | 18 | 41 | 1220 |
| 10 | 6 | 355 | 22 | 22 | 404 | 18 | 6 | 46 | 879 |
| **TOTAL** | 117 | 5766 | 303 | 318 | 6059 | 329 | 126 | 469 | 13487 |

**Table S8D. Tucunacá: Total Size of Structural Variations by Chromosome (bp)**

| Chromosome | Size (Bp) | Contrac-tion | Deletion_ Mobile | Deletion_ Novel | Duplication | Insertion_ Mobile | Insertion_ Novel | Inversion | Translocation | TOTAL | % occupied |
|---|---|---|---|---|---|---|---|---|---|---|---|
| 1 | 127,168,885 | 31,618 | 12,670,731 | 17,701 | 232,374 | 12,659,902 | 15,253 | 355,944 | 244,660,249 | 25,983,523 | 20.43 |
| 2 | 123,336,123 | 12,195 | 11,791,069 | 6,530 | 300,769 | 12,590,062 | 14,889 | 1,197,053 | 228,720,847 | 25,912,567 | 21.01 |

| | | | | | | | | | | | |
|---|---|---|---|---|---|---|---|---|---|---|---|
| 3 | 119,475,968 | 18,975 | 9,740,692 | 9,646 | 40,576 | 12,141,487 | 7,422 | 433,694 | 226,719,622 | 22,392,492 | 18.74 |
| 4 | 117,766,653 | 26,519 | 9,883,336 | 12,346 | 25,504 | 10,275,538 | 8,224 | 450,359 | 232,517,146 | 20,681,826 | 17.56 |
| 5 | 115,567,930 | 2,847 | 10,382,352 | 9,041 | 190,787 | 10,770,158 | 8,256 | 1,064,364 | 213,060,374 | 22,427,805 | 19.41 |
| 6 | 113,259,395 | 939 | 10,679,227 | 4,776 | 132,887 | 11,544,502 | 10,194 | 1,797,011 | 206,606,786 | 24,169,536 | 21.34 |
| 7 | 108,960,230 | 167,994 | 8,061,849 | 6,748 | 15,154 | 8,343,968 | 9,989 | 944,381 | 206,004,603 | 17,550,083 | 16.11 |
| 8 | 105,462,586 | 51,556 | 9,679,779 | 7,708 | 38,642 | 10,238,237 | 6,347 | 1,575,783 | 205,217,801 | 21,598,052 | 20.48 |
| 9 | 104,826,184 | 26,355 | 8,026,560 | 10,441 | 28,617 | 9,098,726 | 13,620 | 3,016,211 | 197,344,711 | 20,220,530 | 19.29 |
| 10 | 78,383,799 | 2,800 | 5,618,378 | 4,775 | 26,328 | 6,924,548 | 4,467 | 632,891 | 152,227,324 | 13,214,187 | 16.86 |
| **TOTAL** | 1,114,207,753 | 341,798 | 96,533,973 | 89,712 | 1,031,638 | 104,587,128 | 98,661 | 11,467,691 | 2,113,079,463 | 214,150,60 1 | 19.22 |
| **% occupied** | | 0.03 | 8.66 | 0.01 | 0.09 | 9.39 | 0.01 | 1.03 | 189.65 | 19.22 | |

**Table S9. Repeat annotation of the analysed Malpighiales genomes**

Repeat classes, superfamilies, element counts, cumulative length, and genome occupancy estimated for the *Banisteropsis caapi* var. caupuri (A) and tucunacá (B) and Malpighia emarginata (C) genomes.

***Provided as an external Excel file***

**Table S10. Ka/Ks estimates for duplicated gene pairs in caupuri (A) and tucunacá (B)**

Pairwise Ka, Ks, Ka/Ks, and associated test statistics for duplicated gene pairs identified in the genomes. Ka/Ks was reported as NA when Ks was equal to zero or when a valid ratio could not be calculated.

***Provided as an external Excel file***

**Table S11. LARD-associated genes across caupuri, tucunacá, and acerola (*M. emarginata*)**

Gene-centric catalog of loci associated with LARD annotations, including overlap class, genomic coordinates, representative LARD interval, functional annotation, orthogroup assignment, and CAFE5 status.

***Provided as an external Excel file***

**Table S12. OrthoFinder orthogroups inferred from chromosome-anchored gene sets across the comparative species dataset**

Gene membership is reported for each species. Orthogroups represented only in the caupuri or tucunacá assembly are identified in the *B. caapi* OG status column. Representative annotations for assembly-restricted orthogroups were obtained by mapping gene-level functional descriptions from Table S6A and S6B to the corresponding orthogroup members; the number of genes supporting each representative annotation is also reported.

***Provided as an external Excel file***

**Table S13. Prioritized candidate orthogroups associated with *B. caapi* morphotype divergence**

(A) Candidate orthogroups prioritized from reconstructed terminal CAFE5 events in the chromosome-restricted caupuri and tucunacá gene sets. Candidate OGs were selected using GO-driven prioritization, with Pfam domains as supporting evidence and keyword matches only as fallback evidence. Functional-category columns report evidence-hit counts; zero indicates no support for that category. Branch-specific probability values and significance flags are provided in Table S13. (B) Functional category summary of candidate orthogroups prioritized from reconstructed terminal CAFE5 events.

***Provided as an external Excel file***

**Table S14. CAFE5 expansion and contraction analysis summary**

Gene-level summary of reconstructed terminal CAFE5 gene-family changes in caupuri and tucunacá, including orthogroup identifiers, branch, condition, reconstructed change, branch-specific probability values, significance flags, gene identifiers, and associated functional annotation fields.

***Provided as an external Excel file***

**Table S15. OrthoVenn3 clusters detected only in the caupuri or tucunacá chromosome-level protein sets, integrated with shared-cluster context, OrthoFinder orthogroups, CAFE5 copy-number changes, and functional annotations**

Each row represents an OrthoVenn3 cluster detected only in one sampled genome. Shared OrthoVenn3 clusters assigned to the same OrthoFinder orthogroups are reported as gene-family context. The retained_integrated_candidate field identifies functionally informative exclusive clusters showing directionally concordant overlap with prioritized CAFE5 orthogroups. All exclusive clusters are retained in the table, including clusters excluded from candidate interpretation because of transposable element-related, organellar, or uninformative annotations.

***Provided as an external Excel file***

**Table S16. Candidate-related tandem arrays and adjacent tandem pairs in *B. caapi***

Tandem arrays and candidate-related adjacent tandem pairs overlapping orthogroups prioritized as potentially associated with morphotype divergence.

***Provided as an external Excel file***

**Table S17. Functional enrichment for tandem-duplicated genes in caupuri and tucunacá**

Gene Ontology (GO) enrichment, COG functional category enrichment, and Enzyme Commission (EC) enrichment results for tandem-duplicated genes. For each analysis, genes classified as tandem duplicated and carrying the corresponding annotation type were used as the study set, whereas all genes from the same genome with that annotation type were used as the background population. The columns report the annotation term, term description, morphotype, study and background counts, study and background ratios, fold enrichment, uncorrected P-value, Benjamini–Hochberg FDR, and representative genes supporting each term. Fold enrichment values above 1 indicate overrepresentation among tandem-duplicated genes relative to the annotated genomic background. Terms with FDR $\leq$ 0.05 were considered significant. Representative gene IDs and product annotations are included to facilitate traceability, and complete gene-level support is provided in the accompanying support files.

***Provided as an external Excel file***

**Table S18. Prioritized candidate genes and gene clusters associated with morphotype divergence in *Banisteriopsis caapi***

Candidate loci were selected from significant CAFE5 orthogroups, morphotype-specific OrthoVenn3 clusters, candidate-related tandem arrays, and functional categories associated with development, signaling, transport, hormone response, cell-wall organization, secondary wall or vascular functions, and defense- or stress-related responses. The table reports representative gene IDs or loci, genomic context, orthogroup/cluster/array assignment, functional annotation, main evidence supporting prioritization, candidate rationale, and relative priority. These candidates represent genomic associations with the caupuri and tucunacá morphotypes and do not imply experimentally validated causal effects.

***Provided as an external Excel file***

**Table S19. Genome assemblies used in comparative genomic, phylogenetic, and gene-family analyses**

| Species | Data Source | Version or Accession[a] | Reference |
|---|---|---|---|
| *Banisteriopsis caapi* (caupuri morphotype) | This work | 1.0 | This work |
| *Banisteriopsis caapi* (tucunacá morphotype) | This work | 1.0 | This work |
| *Euphorbia lathyris* | NCBI | GCF_963576675.1 | Wang et al. (2021) [1] |
| *Hevea brasiliensis* | NCBI | GCF_030052815.1 | Cheng et al. (2023) [2] |
| *Malpighia emarginata* | DDBJ[b] | AP035802-AP035862[c] | Shirasawa et al. (2024) [3] |
| *Manihot esculenta* | Phythozome | v8.1 | Bredeson et al. (2016) [4] |
| *Mercurialis annua* | NCBI | GCF_937616625.2 | Christenhusz et al. (2024) [5] |
| *Populus trichocarpa* | Phythozome | v4.1 | Tuskan et al. (2006) [6] |
| *Ricinus communis* | Phythozome | v0.1 | Chan et al. (2010) [7] |
| *Salix purpurea* | Phythozome | v5.1 | Zhou et al. (2020) [8] |
| *Vitis vinifera* | Phythozome | v2.1 | Jaillon et al. (2007) [9] |

[a] Preferably, genomes were downloaded from the Phytozome database. When unavailable, data were obtained from the NCBI Genomes database.

[b] This assembly was only available in the DNA Data Bank of Japan (DDBJ; https://www.ddbj.nig.ac.jp/).

[c] *Malpighia emarginata* genome assembly was reannotated following the same pipeline employed in the *B. caapi* genomes annotation.
